# PROTAC molecule-mediated SpCas9 protein degradation for precise genome editing

**DOI:** 10.1101/2025.01.07.631496

**Authors:** Shengnan Sun, Renhong Sun, Minkang Tan, Maarten Kip, Xiaobao Yang, Baoxu Pang

## Abstract

CRISPR (clustered regularly interspaced short palindromic repeats) technology has revolutionized both fundamental research and genomic medicine. The need for precise control of SpCas9-based CRISPR genome editing has led to the development of various strategies to regulate Cas9 activity, aiming to minimize off-target effects. Proteolysis-targeting chimeras (PROTACs) use the ubiquitin-proteasome system to selectively degrade target proteins. Here, we developed CASPROTAC 6, a reversible and cell-permeable PROTAC degrader that directly targets SpCas9 proteins for degradation. The CASPROTAC 6 molecule was derived by linking a known SpCas9 binder BRD7087 with an S-substituted Lenalidomide cereblon (CRBN) ligand via a six-carbon alkyl linker. Our data show that CASPROTAC 6 is a broad-spectrum degrader of SpCas9 and variants such as dCas9 and Cas9 nickases. CASPROTAC 6 degrades SpCas9 and the SpCas9-guide RNA complex via the proteasome system by approximately 50%. As a result, CASPROTAC 6 effectively enhanced the precision of CRISPR/Cas9-mediated genome editing by increasing target specificity. CASPROTAC 6 may serve as modulators of CRISPR/SpCas9-associated nucleases, enabling precise genome editing while reducing off-target effects and enhancing biosafety.

## Introduction

CRISPR (clustered regularly interspaced short palindromic repeats) technology has revolutionized genome manipulation and opened new venues for both basic research and genomic medicine[1, 2]. CRISPR-associated systems (Cas) were initially discovered as the adaptive immune responses in many bacteria and archaea against bacteriophages[3, 4]. Currently, the most widely used system for genome editing is the CRISPR/Streptococcus pyogenes Cas9 (SpCas9), which introduces double-strand breaks (DSBs) via guide RNA (gRNA). In general, the SpCas9-gRNA complex, recognizes a 20 bp target DNA sequence adjacent to a protospacer adjacent motif (PAM), typically NGG[5]. Then, the complex will induce a DNA double-strand break at 3 bp upstream of the PAM sequence. These DSBs are repaired through either homologous recombination or nonhomologous end joining (NHEJ), the latter of which can result in imprecise repair or insertions, thereby introducing permanent genomic changes[6]. Based on the CRISPR/SpCas9 system, the CRISPR/dCas9 (catalytic inactive or “dead” SpCas9) has also been developed for genome editing. It retains DNA-binding ability via gRNA but lacks nuclease activity. When fused to effector domains—such as VP64 for gene activation, KRAB for repression, or base editors for nucleotide conversions—CRISPR/dCas9 enables epigenomic modulation or permanent genome editing without inducing DSBs[7].

Although the CRISPR/SpCas9 systems are easy to use, potential off-target effects remain a major concern for both basic research and clinical applications[8, 9]. Recent evidence suggests that even on-target activity of CRISPR/SpCas9 could cause unintended genome editing in some cases, such as chromosomal loss within human embryo cells[10], probably due to prolonged on-target editing. To mitigate off-target DNA cleavage, anti-CRISPR proteins (Acrs) have been genetically fused to SpCas9. These Acr-SpCas9 variants fine-tune the nuclease’s activity, lowering it to desirable levels. As a result, they maintain comparable on-target editing efficiency while reducing off-target events, thereby improving genome editing fidelity[11]. This suggests that off-target effects may stem primarily from the hyperactivity of SpCas9. Small-molecule inhibitors have also been developed to regulate SpCas9 activity[12, 13]. However, both approaches have some limitations. Delivering small inhibitory proteins into cells or organisms remains challenging, and their effectiveness often depends on protein half-life. Small-molecule inhibitors, on the other hand, typically act in a dose-dependent manner and may carry unwanted side effects[14].

Proteolysis-targeting chimeras (PROTACs) are an emerging technology that utilizes the intracellular ubiquitin-proteasome system (UPS) to selectively degrade the protein of interest (POI). PROTAC functions as a chimera molecule that binds to the POI on one end and recruits an E3 enzyme on the other end. By bringing the target protein close to the E3 enzyme, it recruits the E2 enzyme to transfer ubiquitin to the POI. The poly-ubiquitinated POIs are then degraded via the UPS. This targeted degradation strategy may offer a promising approach to control CRISPR/SpCas9 activity. Computational modeling has demonstrated that the ratio of on-target to off-target cleavage decreases over time during genome editing [15-18]. Therefore, reducing the half-life of SpCas9 and its variants may help reduce off-target effects. In this study, we developed a series of PROTAC molecules designed to degrade SpCas9, dCas9, and nCas9 proteins. Through optimization, we identified CASPROTAC 6 as the most effective compound. CASPROTAC 6 degrades SpCas9, dCas9, nCas9, and RNP complexes by approximately 50%, thereby enhancing the specificity of CRISPR-mediated genome editing. This molecule holds potential for genome medicine applications where temporal regulation of SpCas9 and its variant protein levels is critical to improving editing fidelity and minimizing unintended effects.

## Results

### Design of CASPROTAC molecules for SpCas9 protein degradation

As previously shown in computational analyses, prolonged or elevated genome editing activity may exacerbate off-target effects[15-18]. To minimize these effects, controlling the half-life of SpCas9 proteins presents a promising strategy. We first assessed the half-life of SpCas9 and dCas9 proteins and observed that both remained in cells for approximately 5-7 days (Supplementary Figure 1). To model how both on-target editing and off-target effects accumulate over time, we designed GFP-Offtarg gRNAs (with 1 or 2 nucleotide mismatches relative to the gRNA targeting the GFP sequence, i.e. GFP Ontarg) to mimic the off-target effects of CRISPR/SpCas9-mediated genome editing by targeting the same GFP coding sequence. Results showed that the on-target gRNA efficiently edited the GFP sequence, leading to a rapid loss of fluorescence and reaching a plateau (∼47%) by day 3. During the same period, the two GFP-Offtarg gRNAs also edited the GFP sequence and reached their editing plateau on day 3, with an off-target editing efficiency of 16.5% and 1.43%, respectively (Supplementary Figure 2). These results align with previous observations that CRISPR/SpCas9 could recognize and cleave DNA sequences with partial mismatches[19], and that such off-target activity can accumulate as long as SpCas9 remains in the cell.

To reduce the half-life of SpCas9 and mitigate off-target effects, we developed a set of PROTAC molecules that directly target SpCas9 for proteasomal degradation, designated as CASPROTACs (Figure 1). The warhead (WH) of CASPROTACs was derived from BRD7087, a validated SpCas9:gRNA binder confirmed through ^19^F NMR spectroscopy [20]. A series of CASPROTACs was synthesized by connecting this warhead to S-substituted lenalidomide, a cereblon (CRBN) ligand, via alkyl linkers of varying lengths (Figure 1, 2 and Supplementary File 1, 2) [21, 22]. Then, the degradation efficacy of these molecules was tested in K562 cells. K562 cells were electroporated with 60 pmol of recombinant SpCas9 proteins, a dosage commonly used in genome editing experiments [23]. One day post electroporation, cells were treated with 1µM of the respective CASPROTACs (1 - 4), or the warhead alone (as the control) for another 24 hrs, before the level of the recombinant SpCas9 proteins was tested. Western blot results showed that CASPROTAC 3 and CASPROTAC 4, which contain longer linkers, demonstrated superior SpCas9 degradation (Figure 2 B-C). We further tested the optimal concentration of these CASPROTAC molecules for degrading SpCas9 proteins using serial dilutions. These two CASPROTACs achieved the best degradation effect at a concentration of 1 µM, with a maximum degradation (Dmax) of 20% and exhibited a clear “hook effect”, a known pharmacodynamic feature of PROTAC molecules (Figure 2 D-E).

**Figure 1:**
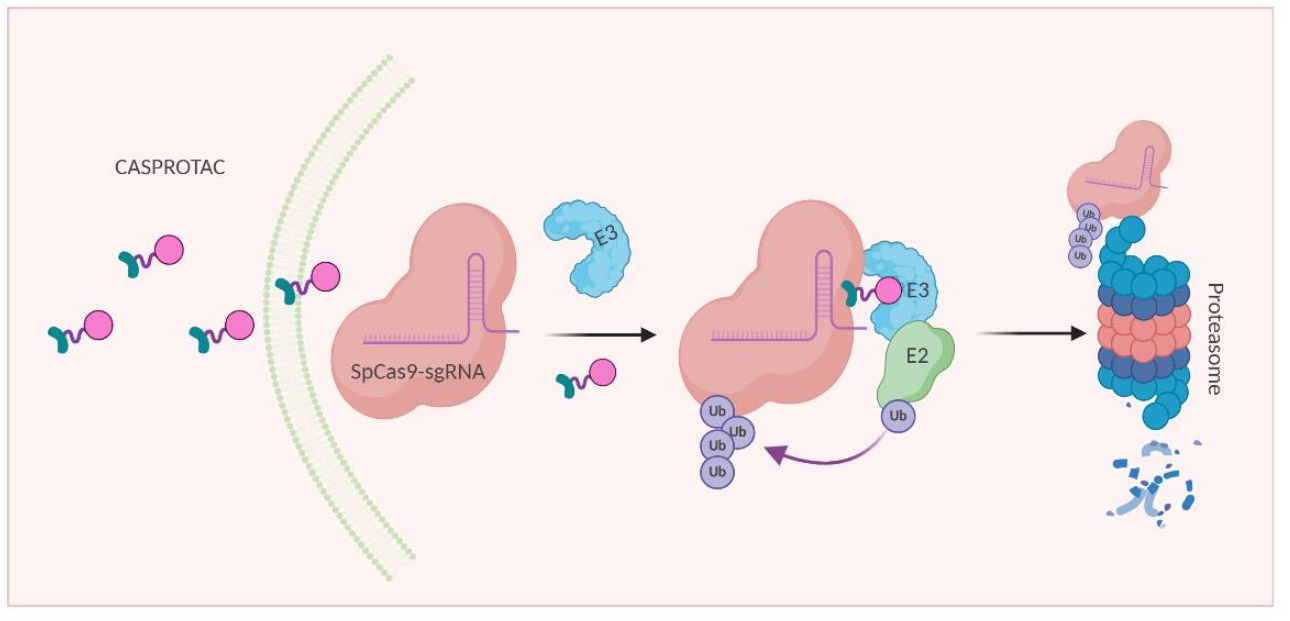
The workflow of CASPROTAC molecules. CASPROTAC works by designing a chimera molecule that binds to the protein of interest on the one end (derived from SpCas9 binder BRD7087) and binds to an E3 enzyme on the other end (S-substituted Lenalidomide, a novel cereblon ligand). Then the E3 enzyme will recruit the E2 enzyme that transfers the ubiquitin to the SpCas9, leading to the degradation of the polyubiquitinated SpCas9 protein.

**Figure 2:**
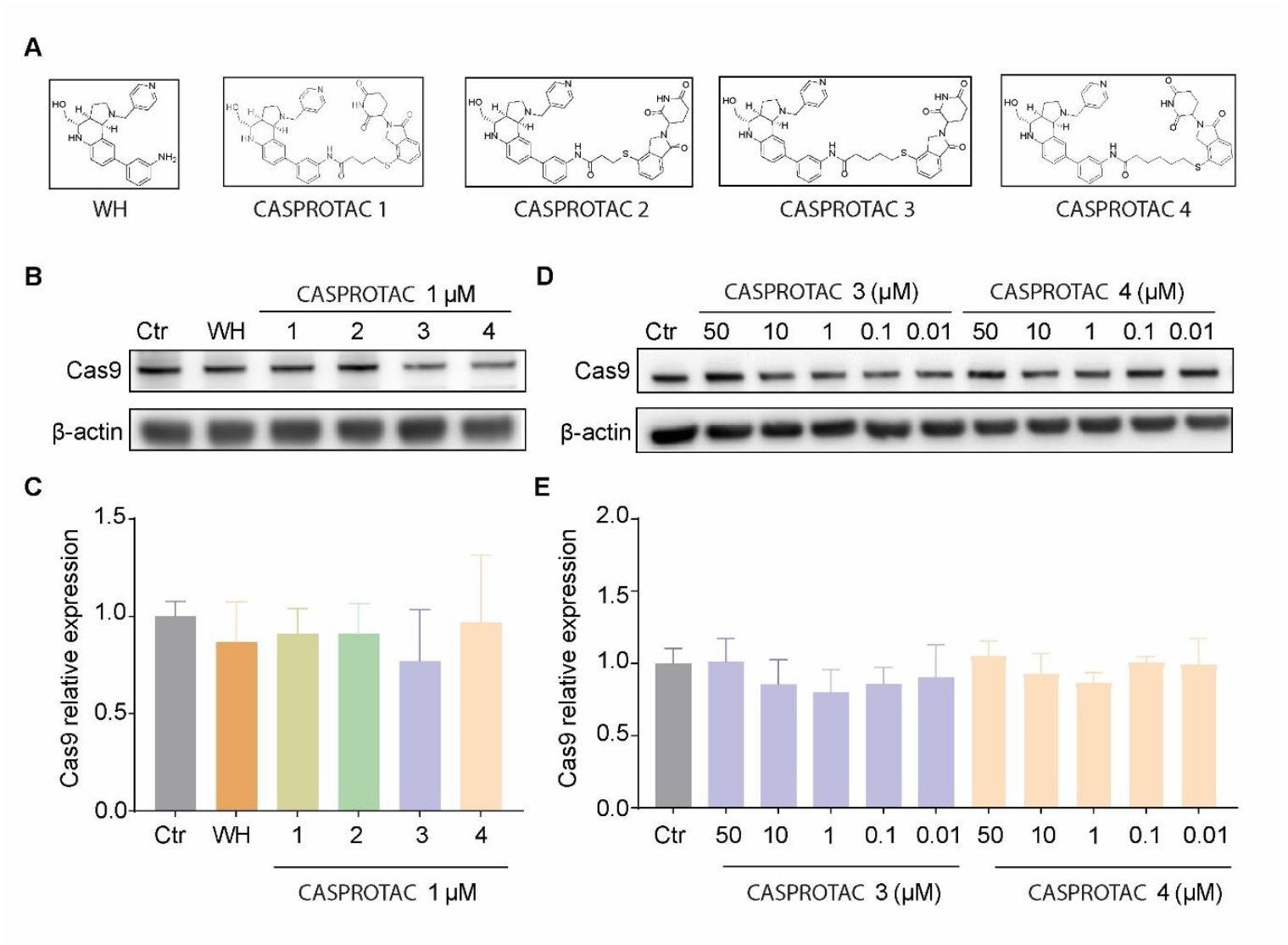
The initial design of CASPROTAC molecules exhibited degradation efficacy. **A**. Chemical structures of CASPROTACs 1-4. **B**. Western blotting was used to test the efficacy of the first batch of CASPROTACs in degrading SpCas9 proteins. K562 cells were electroporated with 60 pmol of SpCas9 protein, followed by incubation with warhead and CASPROTACs 1-4 (1 µM) for 24 hrs. **C**. Quantification of the intensity of the staining from B. **D**. K562 cells were electroporated with 60 pmol of SpCas9 protein, followed by incubation with CASPROTACs 3 and 4 in a dose-dependent manner. Western blotting was used to quantify the efficacy of SpCas9 protein degradation. **E**. Quantification of the intensity of the staining from D. Anti-HA antibody was used to detect the level of recombinant Cas9-HA proteins. Anti- β-actin was used as the loading control. Bars show mean value ± s.e.m. and significance was calculated using Student’s t-test (n = 2 or 3). *p < 0.05, **p < 0.005, and ***p < 0.0001 (versus the control).

### Optimized CASPROTAC molecules degrade SpCas9 protein efficiently

Data from the initial CASPROTACs suggest that longer linkers may enhance SpCas9 degradation efficiency. Therefore, to further optimize the molecules, we synthesized a new series of CASPROTAC molecules (CASPROTACs 5-10) featuring varied linker lengths— either shortened or extended (Figure 3 A and Supplementary File 1, 2). These molecules were tested in K562 cells electroporated with SpCas9 proteins, followed by treatment with 1µM of each CASPROTAC for 24 hrs. Among them, CASPROTAC 6 and CASPROTAC 10 exhibited superior performance, achieving approximately 50% reduction of SpCas9 protein levels (Figure 3B and C). To further evaluate potency, we performed serial dilutions of CASPROTAC 6 and 10. Both compounds showed degradation profiles consistent with typical PROTAC behavior, with maximum degradation observed at 1 μM (Figure 3 D-E). Given its higher efficacy, CASPROTAC 6 was selected for further characterization. To determine the optimal treatment duration for CASPROTAC 6, time-course experiments were conducted. K562 cells electroporated with SpCas9 were treated with 1 μM CASPROTAC 6, and SpCas9 levels were assessed over time. Results showed that maximum degradation (∼50%) was achieved within 24 hours (Figure 3 F-G). These data demonstrate that the optimized CASPROTAC 6 efficiently degrades SpCas9 protein within 24 hours of treatment, making it a promising tool for the control of CRISPR-Cas9 activity.

**Figure 3:**
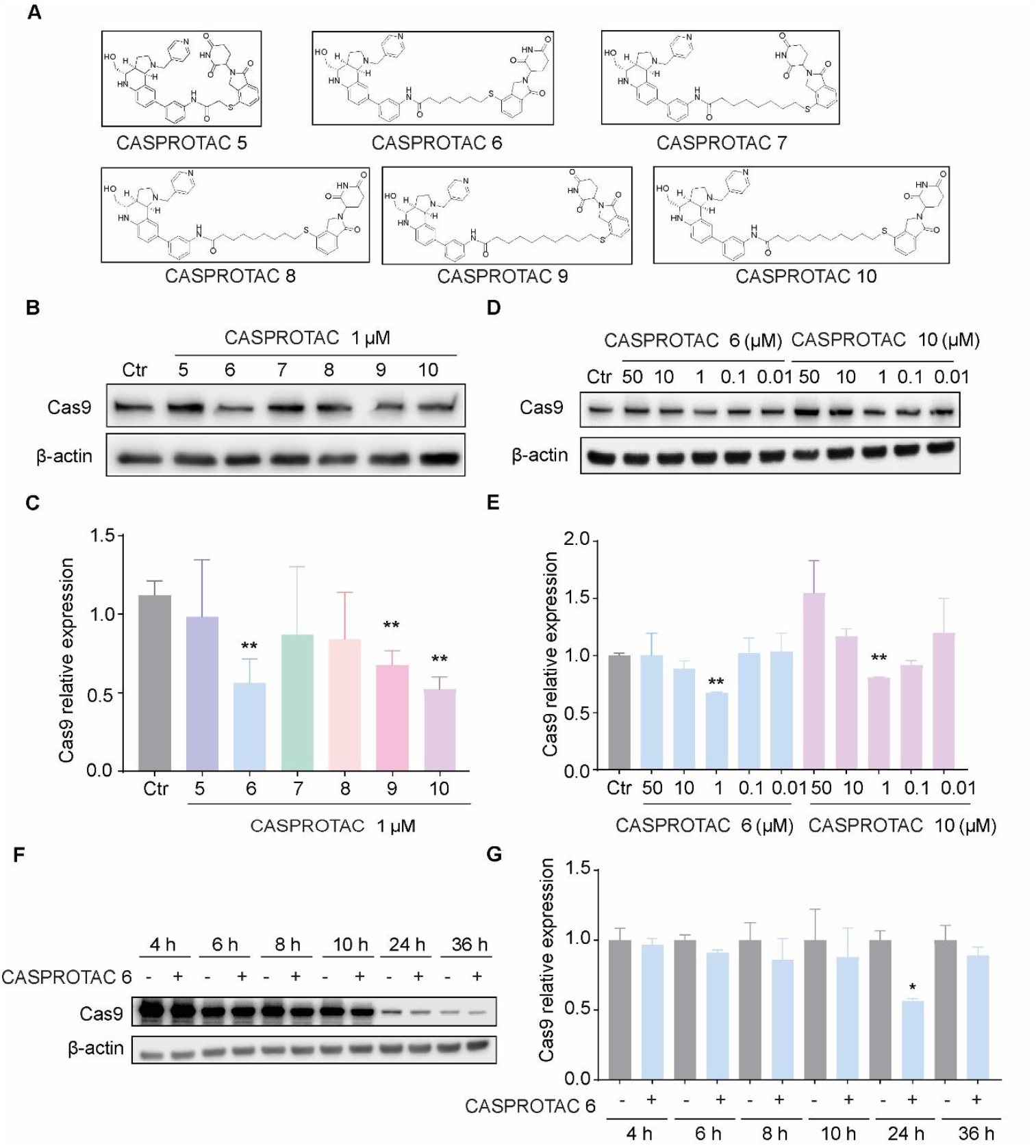
Optimized CASPROTAC molecules degraded SpCas9 efficiently. **A**. Chemical structures of CASPROTACs 5-10. **B**. Western blotting was used to test the efficacy of the optimized CASPROTAC molecules in degrading SpCas9 proteins. K562 cells were electroporated with 60 pmol of SpCas9 protein, followed by incubation with CASPROTACs 5-10 (1 µM) for 24 hrs. **C**. Quantification of the staining intensity from B. **D**. Optimized CASPROTAC molecules resulted in dose-dependent degradation of SpCas9 proteins. Western blotting was used to test the degradation of SpCas9 proteins by CASPROTAC 6 and 10. K562 cells were electroporated with 60 pmol of SpCas9 protein, followed by incubation with 6 and 10 in a dose-dependent manner for 24 hrs. **E**. Quantification of the intensity of the staining from D. **F**. Western blotting was used to test the degradation of SpCas9 proteins by CASPROTAC 6 at different time points. K562 cells were electroporated with 60 pmol of SpCas9 protein were followed at the respective time points with and without CASPROTAC 6 treatment. **G**. Relative quantification was performed between the treated and non-treated cells based on the results from F. Anti-HA antibody was used to detect the level of recombinant Cas9-HA proteins. Anti-β-actin was used as the loading control. Bars show mean value ± s.e.m. and significance was calculated using Student’s t-test (n = 2 or 3). *p < 0.05, **p < 0.005, and ***p < 0.0001 (versus the control).

### The CASPROTAC degrades SpCas9 and the SpCas9-guide-RNA complex via the proteasome system

PROTAC molecules rely on the proteasome degradation system to eliminate the target proteins [24]. To confirm that CASPROTAC 6 functions via this mechanism to degrade SpCas9 proteins, K562 cells electroporated with SpCas9 proteins were co-treated with proteasome inhibitor MG132 (500 nM) and CASPROTAC 6 for 24 hrs. Inhibition of the proteasome completely abrogated SpCas9 degradation, as evidenced by the accumulation of polyubiquitinated proteins in cell lysates (Figure 4 A-C). This result confirms that CASPROTAC 6 operates as a bona fide PROTAC, dependent on proteasomal activity for SpCas9 degradation. Since SpCas9 exerts its function only after assembling with guide RNAs into ribonucleoprotein (RNP) complexes, we further tested whether CASPROTAC 6 is capable of degrading the RNP complexes. The recombinant SpCas9 proteins were pre-incubated with synthesized guide RNAs in vitro to form functional RNP complexes, which were then electroporated into K562 cells and treated with CASPROTAC 6. As shown in Figure 4 D and 4 E, CASPROTAC 6 effectively degraded both free SpCas9 protein and its RNP complexes, indicating its capacity to target multiple functional states of SpCas9. These results underscore the potential of CASPROTAC 6 as a flexible and efficient regulator of CRISPR-based genome editing.

**Figure 4:**
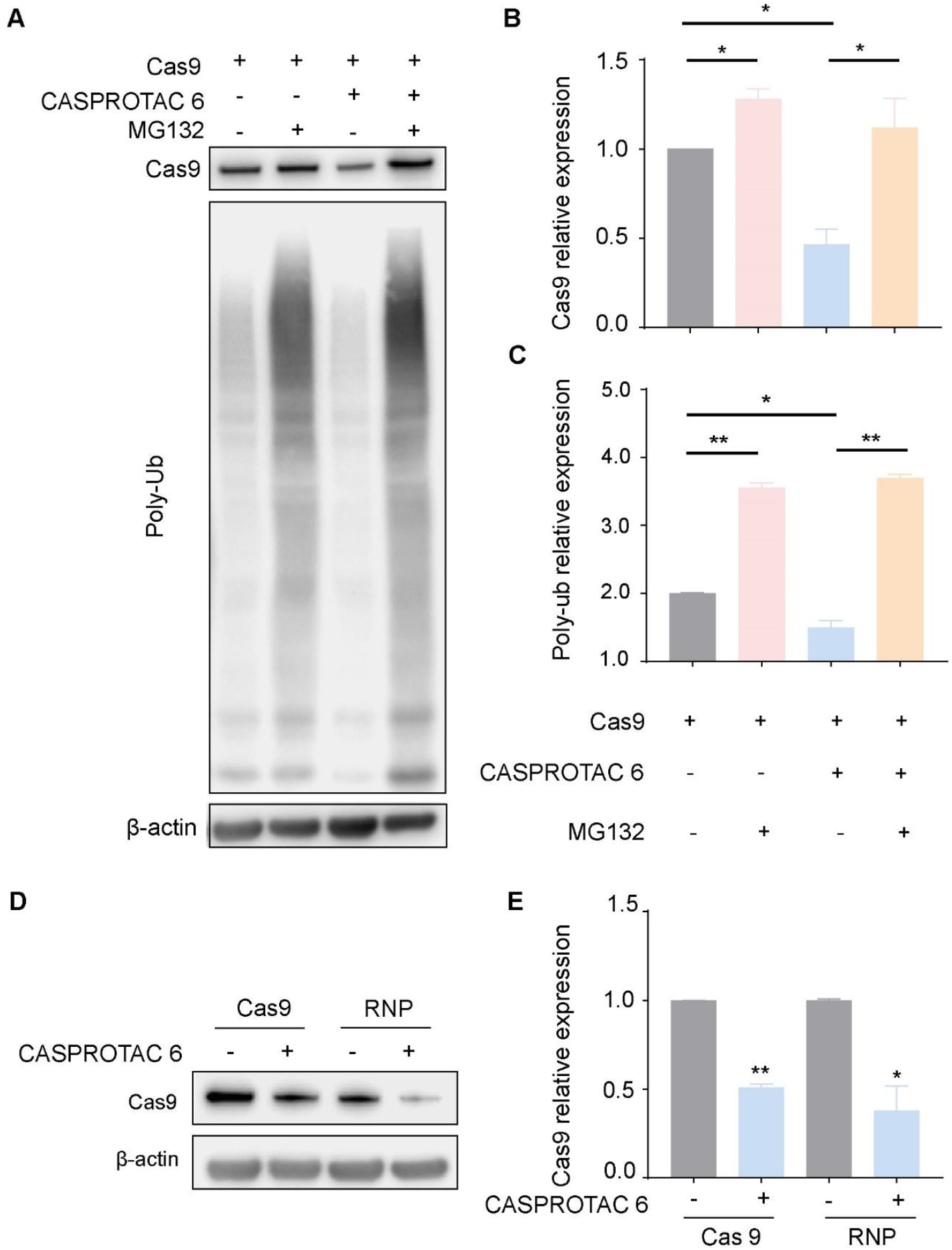
The optimized CASPROTAC molecule degrades SpCas9 and the Cas9-guide-RNA complex via the proteasome system. **A**. The proteasome inhibitor MG132 blocked the degradation of electroporated SpCas9protein by CASPROTAC 6 in K562 cells. Western blotting was used to test CASPROTAC 6 in K562 cells after electroporation with 60 pmol of SpCas9 protein, followed by incubation with CASPROTAC 6 (1 µM) and MG132(500 nM) for 24 hrs. **B**. The intensity of the SpCas9 protein staining was quantified based on A. **C**. The intensity of the poly-ubiquitinated protein staining was quantified based on A. **D**. CASPROTAC 6 degraded CRISPR ribonucleoprotein complexes efficiently. Mix the EGFP-guide RNA and SpCas9 protein components at a 1:1.2 molar ratio (60 pmol SpCas9 + 72 pmol guide RNA-EGFP for 1 million cells). Incubate at room temperature for 20 minutes to allow the formation of the CRISPR ribonucleoprotein complexes. Western blotting for CASPROTAC 6 in K562 cells after electroporation with 60 pmol of SpCas9protein or RNP, followed by incubation with CASPROTAC 6 for 24 hrs. **E**. Quantification of the intensity of the staining from D. Anti-HA antibody was used to detect the level of recombinant Cas9-HA proteins. Anti-β-actin was used as the loading control. An anti-ubiquitin antibody was used to assess the general ubiquitin levels of proteins. Bars show mean value ± s.e.m. and significance was calculated using Student’s t-test (n = 2 or 3). *p < 0.05, **p < 0.005, and ***p < 0.0001 (versus the control).

### CASPROTAC 6 degrades CRISPR/SpCas9 variants and reduces CRISPR editing efficiency

In addition to wild-type SpCas9, which introduces DNA DSBs for genome editing, various engineered SpCas9 variants have also been developed for diverse applications, including generating DNA single-strand breaks or binding DNA without inducing cleavage. The dCas9 protein, for example, binds DNA without introducing DSBs and is widely used in epigenomic modulation and transcriptional regulation. To test if the CASPROTAC 6 can also target and degrade dCas9 proteins, the 293T cells that conditionally express dCas9-APEX2 were treated with CASPROTAC 6. These cells express dSpCas9-APEX2 upon doxycycline treatment, and the expression stops once doxycycline is removed. Using this inducible system to monitor the degradation efficiency of dCas9 proteins by CASPROTAC 6, the results showed that 1 μM CASPROTAC 6 degraded dCas9-APEX2 proteins by 50% within 24 hrs, matching its efficiency against SpCas9 proteins (Figure 5 A-B). Furthermore, similar to the effects on SpCas9 proteins, a dose-dependent degradation of dCas9 proteins was also observed with CASPROTAC 6 (Figure 5 A-B).

**Figure 5:**
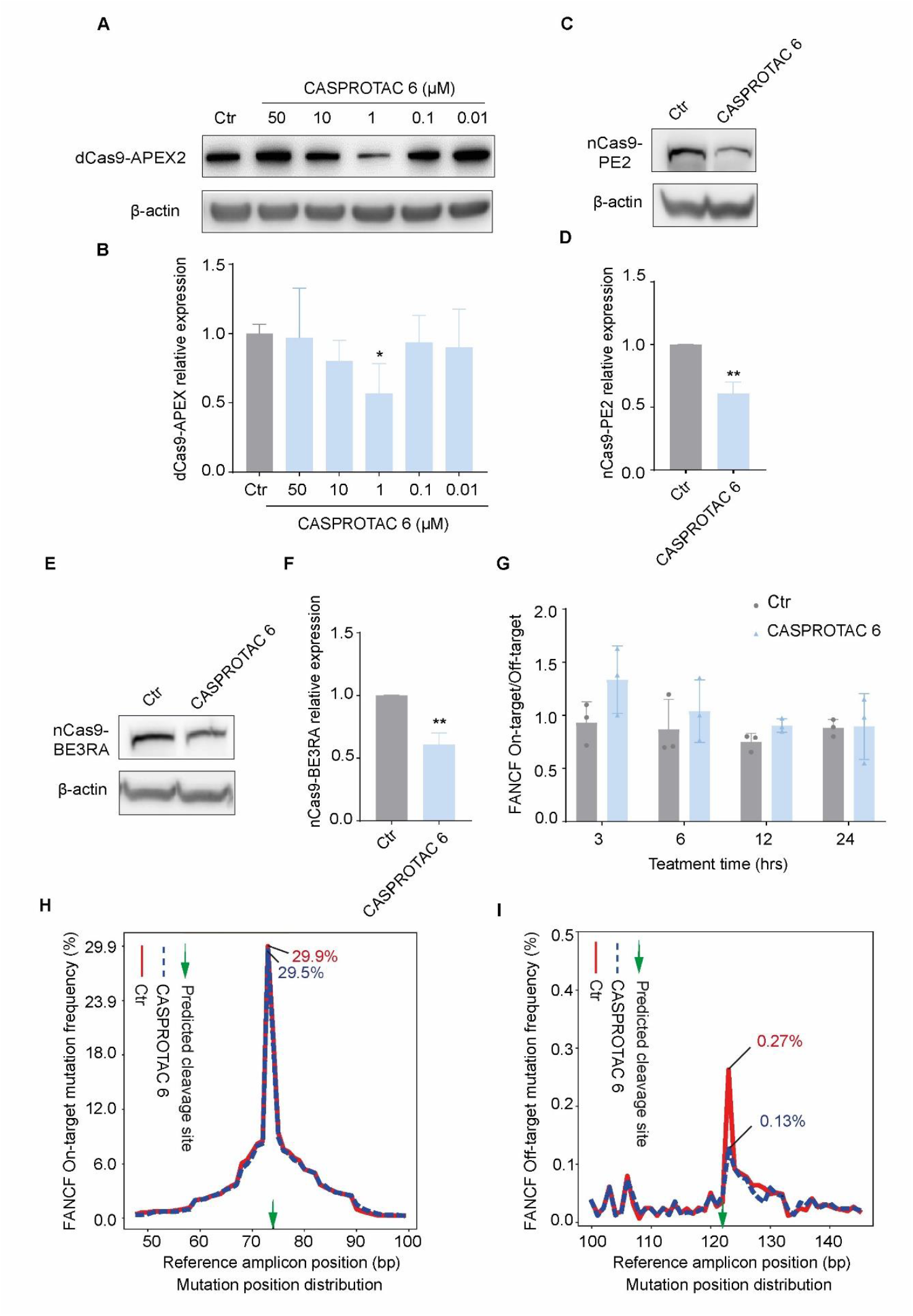
Optimized CASPROTAC 6 degraded CRISPR/SpCas9 toolbox and reduced CRISPR editing efficiency. **A**. Optimized CASPROTAC 6 degraded dCas9 proteins. 293T-Tet-On 3G cells with stable integration of dCas9-APEX2 were pretreated with doxycycline (500 ng/ml) for 9 hrs to the simultaneous expression of Cas9-APEX2 protein complex, 15 hrs after removing doxycycline, CASPROTAC 6 was added for 24 hrs in a dose manner. The protein level of dCas9-APEX2 was monitored by Western blotting. **B**. Quantification of the intensity of the staining from A. Anti-Flag antibody was used to detect the level of dCas9-APEX-Flag proteins. Anti-β-actin was used as the loading control. **C**. The 293T-nCas9-PE2 bulk cells were pretreated with doxycycline (500 ng/ml) for 24 hrs, then doxycycline was removed and CASPROTAC 6 was added for 24 hrs. The protein level of nCas9-PE2 was monitored by Western blotting. Anti- Cas9 antibody was used to detect the level of nCas9-PE2 proteins. Anti-β-actin was used as the loading control. **D**. Quantification of the intensity of the staining from C. **E**. 293T-nCas9-BE3RA bulk cells were pretreated with doxycycline (500 ng/ml) for 24 hrs. Then, doxycycline was removed and CASPROTAC 6 was added for another 24 hrs. The protein level of nCas9-BE3RA was monitored by Western blotting. Anti-Cas9 antibody was used to detect the level of nCas9-BE3RA proteins. Anti-β-actin was used as the loading control. **F**. Quantification of the intensity of the staining from E. **G-I**. CASPROTAC 6 enhances the targeting specificity of CRISPR/SpCas9-mediated genome editing. FANCF-gRNA and SpCas9 protein were mixed at a 1:1 molar ratio (20 pmol SpCas9+ 20 pmol FANCF-gRNA for 1 million cells) and incubated at room temperature for 20 min to form the RNP complexes. K562 cells were electroporated with FANCF-RNP. Six hours post electroporation, cells were treated with CASPROTAC 6 and collected at the indicated time points for analysis. Temporal profiling of normalized on-target to off-target editing ratios of FANCF-RNP-mediated editing, in CASPROTAC 6-treated vs. untreated cells, assessed by amplicon sequencing (G). Representative mutation frequency at the FANCF on-target site (within 1 bp of the predicted cleavage site), comparing CASPROTAC 6-treated and untreated cells at the 3-hour treatment time point (H). Representative mutation frequency at the FANCF off-target site (within 1 bp of the predicted cleavage site), comparing CASPROTAC 6-treated and untreated cells at the 3-hour treatment time point (I). Bars show mean value ± s.e.m., and significance was calculated using Student’s t-test (A-F: n = 2 or 3, G: n=3). *p < 0.05, **p < 0.005, and ***p < 0.0001 (versus the control).

Prime editors (PE) and base editors (BE) were recently developed and widely used genome-editing tools. They utilize SpCas9 nickase (nCas9) fused to specialized enzymes that mediate genome editing[25]. To further investigate whether the CASPROTAC 6 is a broad-spectrum degrader able to degrade nCas9 fused to distinct effector enzymes, 293T cells expressing inducible nCas9-PE2 or nCas9-BE3RA were treated with CASPROTAC 6. The results showed that CASPROTAC 6 also degraded nCas9 with different effector enzymes within 24 hrs at 1 µM concentration (Figure 5 C-F), suggesting that CASPROTAC 6 acts as a broad-spectrum degrader of SpCas9 and its functional variants.

Finally, to directly evaluate the effect of CASPROTAC 6 on CRISPR/SpCas9-mediated editing outcomes, K562 cells were electroporated with the RNP targeting the region Chr:22,625,792-22,625,811 (FANCF site 2) [26]. Six hours after electroporation, the transfected cells were treated with CASPROTAC 6 for the indicated durations. Amplicon sequencing revealed an increased on-target to off-target editing ratio following CASPROTAC 6 treatment (Figure 5 G), indicating improved editing specificity. Quantitative analysis revealed that CASPROTAC 6 treatment did not affect the mutation frequency near the predicted on-target cleavage site of FANCF within the first 3 hours, compared to the control group (Figure 5 H). Notably, at the predicted off-target site of FANCF, CASPROTAC 6 reduced the editing frequency by approximately 50% within 1 bp of the predicted cleavage site (Ctr: 0.27%/CASPROTAC 6: 0.13%) during the same timeframe, highlighting its potential to enhance genome editing specificity (Figure 5I). In summary, CASPROTAC 6 effectively degrades wild-type SpCas9 as well as dCas9 and nCas9-based genome editors, thereby enhancing the precision of CRISPR/Cas9-mediated genome editing.

## Discussion

Genome editing via the CRISPR/SpCas9 system has revolutionized modern genetic research and genome medicine. Despite its power and versatility, the system presents limitations and safety concerns—particularly in therapeutic contexts. While CRISPR/SpCas9 enables precise on-target genome modifications, it can also induce undesirable genotoxic effects by cleaving off-target sites[27]. It is estimated that the human genome contains approximately 2.98 × 10^8^ “NGG” PAM sites, equivalent to one potential SpCas9 targeting site every ∼10 base pairs[28]. In practice, CRISPR/SpCas9 can tolerate 1- or 2-nucleotide mismatches in the gRNA-target DNA region, which increases the likelihood of off-target cleavage[19, 27]. Though these events are less frequent, unwanted off-target editing may still arise. If such off-target editing happens in the coding regions of essential genes, it may lead to severe side effects in genome medicine, highlighting the urgent need for more precise regulation of SpCas9 activity.

High-throughput screening previously identified BRD0539 as a small-molecule SpCas9 inhibitor capable of blocking CRISPR activity[20]. However, its inhibitory effect is reversible due to non-covalent binding. By contrast, PROTACs are heterobifunctional molecules that irreversibly degrade POIs by bridging them to E3 ubiquitin ligases, leading to their proteasomal degradation [29]. In this study, we designed a new class of PROTACs targeting SpCas9, termed CASPROTACs, based on the validated SpCas9 binder BRD7087 (Figure 1). After chemical optimization, we identified CASPROTAC 6, which degraded SpCas9 by ∼50% at 1 μM within 24 hours, with the characteristic “hook effect” typical of PROTACs [30](Figure 3 A-B). In comparison, BRD0539 required concentrations >10 μM to achieve similar inhibition in an eGFP-disruption assay[20], suggesting that CASPROTAC 6 is a more potent regulator of SpCas9. Furthermore, to evaluate the broader utility of CASPROTAC 6, we tested its effect on SpCas9 variants, including catalytically inactive dCas9 and nCas9, which are core components of widely used technologies such as base editing, prime editing, and CRISPR-mediated transcription regulation[29]. CASPROTAC 6 effectively degraded dCas9 and nCas9 (Figure 5), demonstrating its capacity to modulate a range of CRISPR-based systems. Importantly, both wild-type and variant SpCas9 proteins function as RNP complexes, and CASPROTAC 6 was also capable of degrading the SpCas9-gRNA RNP complex (Figure 4 D). These findings suggest that CASPROTAC 6 is a broad-spectrum degrader of SpCas9 and its variants. While partial degradation may reduce overall CRISPR activity, it may improve editing precision, as shown by amplicon sequencing that CASPROTAC 6 increased the on-target to off-target editing ratio (Figure 5 G). Additionally, mutation frequency analysis revealed that CASPROTAC 6 reduced off-target editing by 50% within 1 bp of the predicted cleavage site within 3 hours of treatment, while preserving on-target editing frequency near the predicted cleavage site (Figure 5 H–I). CASPROTAC 6 offers a novel strategy to enhance the specificity of CRISPR/Cas9-mediated genome editing and represents a promising tool for both fundamental research and therapeutic applications where genome-editing fidelity is critical.

## Materials and methods

### Chemical synthesis

Reagents and solvents were purchased from commercial sources without further purification unless otherwise indicated. Starting materials of 10 are synthesized according to the patent WO2019196812. The progress of reactions was monitored by thin-layer chromatography (TLC) and/or LC-MS. The final compounds were purified by prep-HPLC and characterized by ^1^H NMR/^13^C NMR and HRMS. NMR spectra were recorded on Ascend 400 MHz Bruker spectrometer (operating at 400 MHz for 1H NMR and 126 MHz for 13C NMR), chemical shifts were reported in ppm relative to the residual CH_3_OD (3.31 ppm ^1^H) or *d*_*6*_-DMSO (2.50 ppm ^1^H) and coupling constants (*J*) are given in Hz. The detail of the synthesis of the BRD7087 analogue was described in Supplemental File 1. Multiplicities of signals are described as follows: s --- singlet, br. s --- broad singlet, d --- doublet, t --- triplet, m --- multiple. High Resolution Mass spectra were recorded on solan X 70 FT-MS spectrometer.

### Lentivirus production and MOI determination

To generate the pLenti-TRE3G-BE3RA-PGK-Puro (Addgene,110846) virus, 293T cells were cultured in 6-well plates. When the cells reached 50% confluence, they were transfected with a mixture containing 1 µg of pLenti-TRE3G-BE3RA-PGK-Puro plasmid, 0.6 µg of psPAX2, 0.4 µg of pCMV-VSV-G, and 6 µg of PEI (Polyscience, 23966) in 50 µl of serum-free medium. The culture medium was refreshed 8--12 hours post-transfection, and the supernatant containing viral particles was collected on the third day following transfection.

### Cell culture and cell line

K562 cells were cultured in RPMI 1640 (Gibco) supplemented with 10% fetal bovine serum (Biowest), L-Glutamine (Gibco), and 1% Penicillin-Streptomycin (Gibco), at 5% CO_2_ in a humidified 37°C incubator. 293T cells were cultured in DMEM (Gibco) supplemented with 10% fetal bovine serum (Biowest), L-Glutamine (Gibco), and 1% Penicillin-Streptomycin (Gibco). For 293T-Tet-On 3G cells generation, the wild-type 293T cell line was grown in 6 well plates (50% confluency) and co-transfected with 1.0 μg Inducible Caspex expression (Addgene 97421) plasmid and 1.0 μg PiggyBac Transposase vector using 6 µg PEI (Polyscience, 23966) in 50 µl of serum-free medium and selected with 3 μg ml^-1^ Puromycin (Biomol) for 2-3 days. Single cells were seeded in 96-well plates and expanded. For the generation of 293T-nCas9-PE2 bulk cells, wild-type 293T cells were seeded in 6 well plates and cultured to approximately 50% confluency. Cells were co-transfected with 1.0 μg of the Xlone-PE2 plasmid (Addgene #136463) and 1.0 μg of the PiggyBac Transposase vector using 6 μg of PEI (Polysciences, Cat. No. 23966) diluted in 50 μl of serum-free medium. Two days post-transfection, cells were selected with 3 μg/ml Blasticidin (Biomol) for 2--3 days to enrich for stable integrated cells.

For the generation of 293T-nCas9-BE3RA bulk cells, wild-type 293T cells were infected with pLenti-TRE3G-BE3RA-PGK-Puro lentivirus at a multiplicity of infection (MOI) of 0.3. Two days post-infection, cells were selected with 3 μg/ml puromycin (Biomol) for 2--3 days. The selected cells were then transfected with 1.0 μg of the XLone-GFP plasmid (Addgene #96930) and 1.0 μg of the PiggyBac Transposase vector. Two days after transfection, GFP expression was induced with doxycycline, and GFP-positive cells were enriched by sorting as stable integrated cells.

### Reagents

Doxycycline was obtained from Sanbio B.V., and the proteasome inhibitor MG132 was purchased from MedChemExpress (Cat. No. HY-13259). Recombinant Streptococcus pyogenes SpCas9 protein, containing an HA tag, was produced by the Protein Facility of Leiden University Medical Center (LUMC) using the pET28a/SpCas9-Cys plasmid (Addgene #53261).

### Guide RNA Synthesis and ribonucleoprotein complexes formulation

The guide RNAs targeting EGFP sequence were produced by in vitro transcription using the following primer sequences:

Common-scaffold-primer: AAAAGCACCGACTCGGTGCCACTTTTTCAAGTTGATAACGGACTAGCCTTATTTTAACTT GCTATTTCTAGCTCTAAAAC;

gRNA-GFP-ontarg: TTCTAATACGACTCACTATAGGACCAGGATGGGCACCACCCGTTTTAGAGCTAGA;

gRNA-GFP-ontarg #1: TTCTAATACGACTCACTATAGGACCAGGATGGGCACCCCCCGTTTTAGAGCTAGA

gRNA-GFP-ontarg #2: TTCTAATACGACTCACTATAGGACCAGGATGTGTACCACCCGTTTTAGAGCTAGA

FANCF-ontarg: TTCTAATACGACTCACTATAGGCTGCAGAAGGGATTCCATGGTTTTAGAGCTAGA

The gRNA primers were annealed with common-scaffold-primer and made into double-strand DNA using NEBNext High-Fidelity 2X PCR Master Mix: 98 °C for 2 mins, 60 °C for 1 min and 72 °C for 3 min. For each reaction, 25 μl of 2X PCR Master Mix, 5 μl of T7 -Guide-EGFP (10 μM), 5 μl of Common scaffold primer (10 μM) and 15 μl of Nuclease-free water were pooled. The pooled PCR products were further purified using the QIAGEN MinElute column. The gRNAs were transcribed from the DNA template and purified according to the manufacturer’s protocol (Agilent SureGuide gRNA Synthesis Kit, 5190-7714, 5190-7719).

To assemble ribonucleoprotein (RNP) complexes, Cas9 protein and sgRNA were mixed at the indicated ratios and preincubated at room temperature for 20 minutes. Following incubation, the RNP complexes were used immediately for downstream applications.

### Western blot

Treated cells were washed with cold phosphate-buffered saline (PBS) and then lysed in 1X RIPA buffer (Thermo Scientific) supplemented with a protease inhibitor cocktail (Roche). The cell lysate was sonicated followed by centrifugation. Protein concentrations were determined using the Pierce BCA Protein Assay Kit (Thermo Scientific). Equal amounts of protein were separated via 8% or 12% SDS-PAGE. The separated proteins were transferred to PVDF membranes, blocked with 5% nonfat milk in PBST for 1 h at room temperature. Membranes were probed with primary antibodies against HA (Cell Signaling Technology H9658, 1:1000), Flag (Sigma F1804, 1:1000), SpCas9 (Cell Signaling Technology 14697S, 1:1000), ubiquitin (Abcam ab19247, 1:1000), and β-actin (Sigma A5441, 1:5000) as the loading control overnight at 4°C. Then, the membranes were washed with PBST three times, incubated with the appropriate secondary antibodies (1:10,000 dilution) for 1 hour at room temperature, and then washed with PBST three times. The resulting signal was visualized using a Bio-Rad ChemiDoc Imaging System with Pierce enhanced chemiluminescence (ECL) Plus Western Blotting Substrate (Thermo Scientific). The intensity of the staining was quantified using Image Lab software (6.0.1).

### Amplicon Sequencing and analysis

K562 cells were electroporated with FANCF-RNP using the SF Cell Line 4D-Nucleofector kit (Lonza) following the pulse program of FF-120. Transfected cells were plated in the T25 flask (Corning), 6 hrs after electroporation, CASPROTAC 6 was added for different durations. The cells were harvested, and the genomic DNA was extracted using the DNeasy Blood & Tissue Kit (Qiagen). Amplicon sequencing samples were prepared via two-step PCR following the protocol reported previously[26]. Amplicon sequences were analyzed by CRISPResso 2.

### Statistical analyses

Data were analyzed using GraphPad Prism software (version 10.2.3). Statistical tests were conducted with the Student’s t-test. P values less than 0.05 were regarded as indicating a statistically significant difference.

## Contributions

B.P. conceived and supervised the study. X.Y. and R.S. designed and synthesized the CASPROTACs. S.S. designed and performed the experiments, and analyzed the data, with the help of M.K. M.T. analyzed the amplicon sequencing data. S.S. and B.P. wrote the manuscript with inputs from all authors.

