## Supplemental File 1 for "PROTAC molecule-mediated SpCas9 protein degradation for precise genome editing"

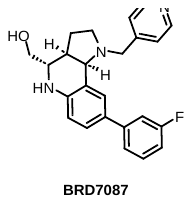

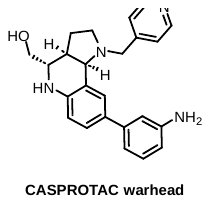

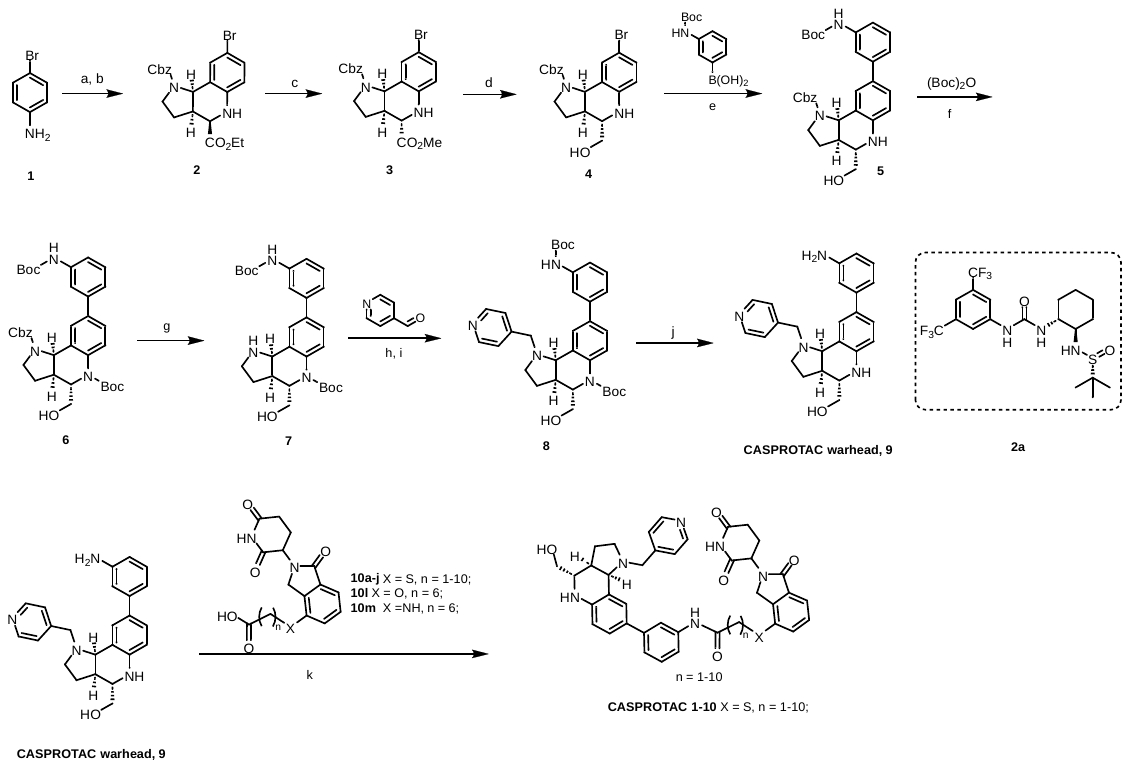

Scheme 1. Synthetic routes for NAMPT PROTAC degraders

Reagents and conditions: (a) ethyl 2-oxoacetate, toluene, MgSO_4_, rt, 1 h; (b) **2a**, benzyl 2,3-dihydro-1H-pyrrole-1-carboxylate, *p*TSA, toluene, 5Å MS, -65 °C, 1 h; (c) NaOMe, MeOH, 50 °C, 0.5 h; (d) LiBH_4_, THF, rt, 3 h; (e) XPhos Pd G3; Cs_2_CO_3_, THF/H_2_O, 100 °C, 1 h; (f) (Boc)_2_O, Et_3_N, DMAP, DCM, rt, 0.5 h; (g) 10% Pd/C, MeOH, H_2_ (50 psi), rt, 2 h; (h) AcOH, DCM, rt, 1 h; (i) NaBH(OAc)_3_, rt, 3 h; (j) 4M HCl/MeOH, MeOH, rt, 3 h; (k) HOAT, EDCI, DIEA, DMF, rt, 1h.
