## Supplemental File 2 for "PROTAC molecule-mediated SpCas9 protein degradation for precise genome editing"

**Chemical synthesis.**

General methods

Reagents and solvents were purchased from commercial sources without further purification, unless otherwise indicated. Starting materials of 10 are synthesized according to the patent WO2019196812. The progress of reactions was monitored by thin-layer chromatography (TLC) and/or LC-MS. The final compounds were purified by prep-HPLC and characterized by ^1^H NMR/^13^C NMR and HRMS. NMR spectra were recorded on Ascend 400 MHz Bruker spectrometer (operating at 400 MHz for 1H NMR and 126 MHz for 13C NMR), chemical shifts were reported in ppm relative to the residual CH_3_OD (3.31 ppm 1H) or *d_6_*-DMSO (2.50 ppm 1H) and coupling constants (J) are given in Hz. Multiplicities of signals are described as follows: s --- singlet, br. s --- broad singlet, d --- doublet, t --- triplet, m --- multiple. High Resolution Mass spectra were recorded on solan X 70 FT-MS spectrometer.

**Scheme S1**. Synthesis of **BRD7087** analogue.

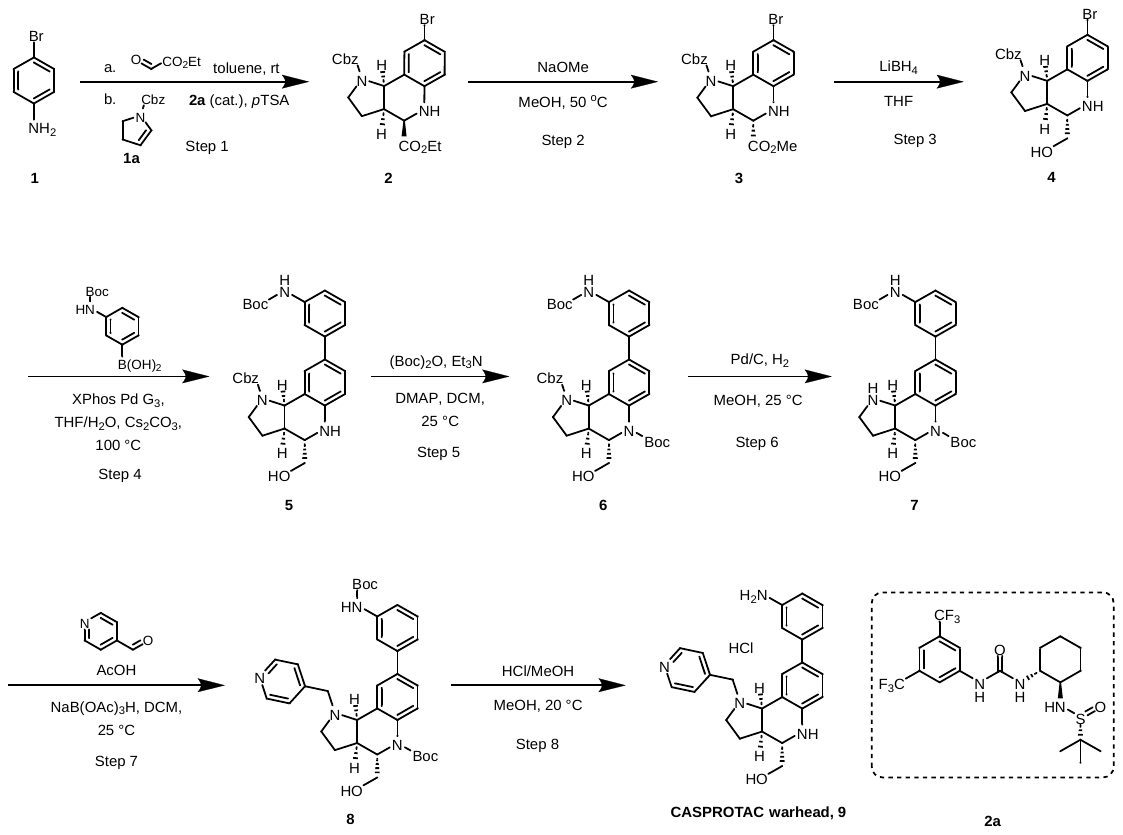

**Step 1**: Preparation of 1-benzyl 4-ethyl (*3*a*R*,*4R*,*9bR*)-8-bromo-2,3,3a,4,5,9b-hexahydro-1*H*-pyrrolo[3,2-*c*]quinoline-1,4-dicarboxylate (compound **2**)

To a solution of ethyl 2-oxoacetate (9.27 g, 45.42 mmol, 50% purity, 1.2 eq) in toluene (40 mL) was added MgSO_4_ (273.35 mg, 2.27 mmol, 0.06 eq). The mixture was cooled to 0 ºC and compound **1** (6.51 g, 37.85 mmol, 1 eq) was added as a solution in toluene (40 mL) over 5 min. The reaction mixture was stirred for 1 h and then MgSO_4_ was filtered off and excess of solvent was removed in vacuo. The resulting product was co-evaporated and then the crude imine was used directly in the next step without further purification. A solution of the crude imine (E)-ethyl 2-((4-bromophenyl)imino)acetate (1 equiv), compound **2a** (358.41 mg, 756.98 umol, 0.02 eq) and compound **1a** (10 g, 49.20 mmol, 1.3 eq) was stirred at 15 ℃ for 5 min in toluene (200 mL) in the presence of 5Å MS (60 mg) before cooling to -65 °C. A solution of anhydrous *p*TSA (65.18 mg, 378.49 umol, 0.01 eq) in THF (20 mL) was added slowly and the reaction mixture was stirred for 1 h. TLC indicated the reaction was complete and one major new spot with lower polarity was detected. The reaction solvent was removed in vacuo and the residue was purified by column chromatography (SiO_2_, Petroleum ether/Ethyl acetate = 10/1 to 5/1) to afford the desired compound **2** as a yellow oil (11 g, 22.31 mmol, 58.94% yield). LC-MS (ESI) m/z: calcd for C_22_H_24_BrN_2_O_4_^+^ [M+H]^+^, 460.35; found, 459.1/461.1.

**Step 2**: Preparation of 1-benzyl 4-methyl (3a*R*,4*S*,9b*R*)-8-bromo-2,3,3a,4,5,9b-hexahydro-1*H*-pyrrolo[3,2-*c*]quinoline-1,4-dicarboxylate (compound **3**)

Compound **2** (11.0 g, 20.48 mmol, 85.53% purity, 1 eq) was dissolved in MeOH (200 mL) under an atmosphere of N_2_. NaOMe (0.1 M, 491.58 mL, 2.4 eq) was added dropwise at 25 °C. The reaction mixture was stirred until complete consumption of starting material was observed (~5 min) to obtain the desired methylester. The reaction mixture was warmed to 50 °C and stirred for 30 min. TLC showed two major new spots with lower polarity were detected. The mixture was quenched with sat. NH_4_Cl (700 mL), and then extracted with Ethyl acetate 600 mL (200mL × 3). The combined organic layers were dried over Na_2_SO_4_, filtered and concentrated under reduced pressure to give a residue. The residue was purified by column chromatography (SiO_2_, Petroleum ether/Ethyl acetate = 5/1 to 3/1) to afford the desired compound **3** as a light yellow oil (3 g, 6.74 mmol, 32.89% yield).

**Step 3**: Preparation of benzyl (3a*R*,4*S*,9b*R*)-8-bromo-4-(hydroxymethyl)-2,3,3a,4,5,9b-hexahydro-1*H*-pyrrolo[3,2-*c*]quinoline-1-carboxylate (compound **4**)

To a solution of compound **3** (3.00 g, 6.74 mmol, 1 eq) in THF (30.0 mL) was added LiBH_4_ (220.13 mg, 10.11 mmol, 1.50 eq) at 0°C. The mixture was stirred at 25 °C for 3 hr. LC-MS showed ~88.59% of desired compound was detected. The reaction mixture was quenched with saturated solution of NH_4_Cl at 0°C, then extracted with Ethyl acetate (200 mL × 3). The combined organic layers were washed with brine (100 mL), dried over Na_2_SO_4_, filtered and concentrated under reduced pressure to give the crude product of compound **4**, which was used to the next step directly as a yellow solid (2.47 g, crude). LC-MS(ESI) m/z: calcd for C_20_H_22_BrN_2_O_3_^+^ [M+H]^+^, 418.31; found, 417.0/419.0.

**Step 4**: benzyl (3a*R*,4*S*,9b*R*)-8-(3-((tert-butoxycarbonyl)amino)phenyl)-4-(hydroxymethyl)-2,3,3a,4,5,9b-hexahydro-1*H*-pyrrolo[3,2-*c*]quinoline-1-carboxylate (compound **5**)

A solution of compound **4** (2.47 g, 5.92 mmol, 1 eq), [3-(tert-butoxycarbonylamino)phenyl]boronic acid (3.51 g, 14.80 mmol, 2.5 eq), XPhos Pd G_3_ (501.02 mg, 591.91 umol, 0.1 eq) and Cs_2_CO_3_ (5.79 g, 17.76 mmol, 3 eq) in THF (45 mL) and H_2_O (9 mL) were stirred at 100 °C for 1 h under N_2_. LC-MS showed ~44.98% of desired compound was detected. The mixture was filtered on a celite pad and then extracted with Ethyl acetate (100 mL × 2). The combined organic phases were dried over Na_2_SO_4_, filtered and concentrated under reduced pressure to give the crude product of comound **5**, which was used to the next step directly as a yellow solid (3 g, crude). LC-MS(ESI) m/z: calcd for C_27_H_28_N_3_O_5_^+^ [M+H-*^t^*Bu]^+^, 474.54; found, 474.4.

**Step 5**: Preparation of 1-benzyl 5-(tert-butyl) (3a*S*,4*S*,9b*R*)-8-(3-((tert-butoxycarbonyl)amino)phenyl)-4-(hydroxymethyl)-2,3,3a,9b-tetrahydro-1*H*-pyrrolo[3,2-*c*]quinoline-1,5(4*H*)-dicarboxylate (comound **6**)

A mixture of compound **5** (3 g, 5.66 mmol, 1 eq) , (Boc)_2_O (1.36 g, 6.23 mmol, 1.43 mL, 1.1 eq) , Et_3_N (687.81 mg, 6.80 mmol, 946.09 uL, 1.2 eq) and DMAP (69.20 mg, 566.44 umol, 0.1 eq) in DCM (30 mL) was stirred at 25 °C for 0.5 hr under N_2_ atmosphere. LC-MS showed ~41.11% of desired compound was detected. The reaction mixture was quenched with water 40 mL and extracted with DCM (50 mL × 3). The combined organic layers were dried over Na_2_SO_4_, filtered and concentrated under reduced pressure to give a residue. The residue was purified by column chromatography (SiO_2_, Petroleum ether/Ethyl acetate= 3/1) to afford compound **6** as a yellow solid (2.5 g, 3.97 mmol, 65.79% yield). LC-MS(ESI) m/z: calcd for C_36_H_43_N_3_O_7_^+^ [M+H]^+^, 629.75; found, 630.5.

**Step 6**: Preparation of tert-butyl (3a*S*,4*S*,9b*R*)-8-(3-((tert-butoxycarbonyl)amino)phenyl)-4-(hydroxymethyl)-1,2,3,3a,4,9b-hexahydro-5*H*-pyrrolo[3,2-*c*]quinoline-5-carboxylate (compound **7**)

A mixture of compound **6** (2.5 g, 3.97 mmol, 1 eq), Pd/C (250 mg, 10% purity) in MeOH (25 mL) was degassed and purged with H_2_ for 3 times, and then the mixture was stirred at 25 °C for 2 hr under H_2_ atmosphere at 50 psi. LC-MS showed ~83.56% of desired compound was detected. The mixture was filtered on a celite pad and then the filtrate was concentrated under reduced pressure to give a crude product of compound **7**, which was used to the next step as a white solid (2 g, crude). LC-MS(ESI) m/z: calcd for C_28_H_37_N_3_O_5_^+^ [M+H]^+^, 495.62; found, 496.5.

**Step 7**: Preparation of tert-butyl (3a*S*,4*S*,9b*R*)-8-(3-((tert-butoxycarbonyl)amino)phenyl)-4-(hydroxymethyl)-1-(pyridin-4-ylmethyl)-1,2,3,3a,4,9b-hexahydro-5*H*-pyrrolo[3,2-*c*]quinoline-5-carboxylate (compound **8**)

A mixture of compound **7** (1.9 g, 3.83 mmol, 1 eq), pyridine-4-carbaldehyde (410.62 mg, 3.83 mmol, 360.20 uL, 1 eq) and AcOH (230.21 mg, 3.83 mmol, 219.25 uL, 1 eq) in DCM (38 mL) was stirred at 25 °C for 1 hr, then to the solution was added NaBH(OAc)_3_ (2.44 g, 11.50 mmol, 3 eq) and the mixture was stirred at 25 °C for another 3 hr. LC-MS showed ~79.80% of desired compound was detected. The residue was diluted with 30 mL DCM, quenched with a saturate NaHCO_3_ aqueous solution (50 mL), then was extracted with DCM (30 mL × 3). The combined organic layers were washed with brine, dried over Na_2_SO_4_, filtered and concentrated under reduced pressure to give a residue. The residue was purified by prep-HPLC with solvent A (0.05% NH_3_·H_2_O in H_2_O) and solvent B (CH_3_CN) as eluents to afford the desired compound **8** as a white solid (1.1 g, 1.87 mmol, 48.90% yield). LC-MS(ESI) m/z: calcd for C_34_H_42_N_4_O_5_^+^ [M+H]^+^, 586.73; found, 587.4.

**Step 8**: Preparation of ((3a*R*,4*S*,9b*R*)-8-(3-aminophenyl)-1-(pyridin-4-ylmethyl)-2,3,3a,4,5,9b-hexahydro-1*H*-pyrrolo[3,2-*c*]quinolin-4-yl)methanol (comopound **9**)

Compound **8** (1.1 g, 1.87 mmol, 1 eq) was dissolved in MeOH (11 mL), then HCl/MeOH (4 M) was added dropwise at 25 °C, and then the mixture was stirred for 3 hr. LC-MS showed ~99.10% of desired compound was detected. The reaction mixture was concentrated under reduced pressure to give the desired compound **9** as a yellow solid (97.30% HPLC purity, 1.1 g, 1.82 mmol, 96.94% yield). ^1^H NMR (400 MHz, D_2_O) *δ* 8.67 (br d, *J* = 5.50 Hz, 2H), 8.03 (br d, *J* = 3.38 Hz, 2H), 7.57 (br d, *J* = 7.38 Hz, 3H), 7.50 (s, 1H),7.45 - 7.35 (m, 1H), 7.31 (br d, *J* = 7.13 Hz, 1H), 6.94 (d, *J* = 8.63 Hz, 1H), 5.03 - 4.87 (m, 2H), 4.81 (br s, 1H), 3.77 (dd, *J* = 11.82, 3.31 Hz, 1H), 3.70 - 3.52 (m, 3H), 3.33 (dt, *J* = 5.44, 2.53 Hz, 1H), 2.84 - 2.73 (m, 1H), 2.59 (br dd, *J* = 14.38, 6.63 Hz, 1H), 2.22 (br s, 1H). LC-MS(ESI) m/z: calcd for C_24_H_26_N_4_O^+^ [M+H]^+^, 386.50; found, 387.3.

**Scheme S2**. Synthesis of **CASPROTAC** **1-10**.

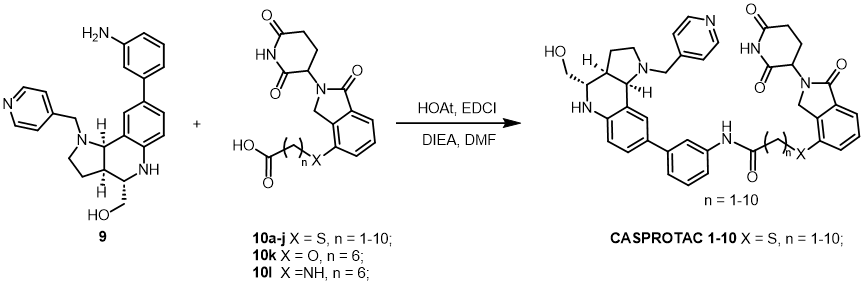

General procedure

To a mixture of compound **9** (10 mg, 0.026 mmol, 1.0 eq),  acid **10** (0.026 mmol, 1.0 eq), EDCI (9.97 mg, 0.052 mmol, 2.0 eq), HOAt (7.08 mg, 0.052 mmol, 2.0 eq) and DMF (1.5 mL) were added DIEA (21.485 μL, 0.130 mmol, 5.0 eq). The mixture was stirred for 1 hour at room-temperature. LC-MS showed the reaction was complete and the major desired product was detected.

**CASPROTAC 1**

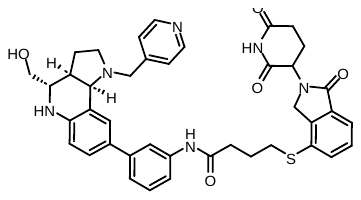

4-((2-(2,6-dioxopiperidin-3-yl)-1-oxoisoindolin-4-yl)thio)-*N*-(3-((3a*R*,4*S*,9b*R*)-4-(hydroxymethyl)-1-(pyridin-4-ylmethyl)-2,3,3a,4,5,9b-hexahydro-1*H*-pyrrolo[3,2-*c*]quinolin-8-yl)phenyl)butanamide (**11c**, 6.5 mg, 34.2% yield, yellow solid). ^1^H NMR (400 MHz, *d_6_-*DMSO) δ 10.99 (s, 1H), 10.06 (s, 1H), 8.76 (d, *J* = 5.9 Hz, 2H), 7.90 – 7.84 (m, 3H), 7.68 – 7.64 (m, 1H), 7.57 (d, *J* = 6.7 Hz, 1H), 7.55 – 7.50 (m, 1H), 7.47 – 7.38 (m, 3H), 7.32 – 7.24 (m, 2H), 6.89 (d, *J* = 8.5 Hz, 1H), 5.12 (dd, *J* = 13.3, 5.1 Hz, 1H), 4.85 (d, *J* = 13.2 Hz, 1H), 4.74 – 4.57 (m, 1H), 4.36 (d, *J* = 17.3 Hz, 1H), 4.22 (d, *J* = 17.3 Hz, 1H), 3.68 – 3.62 (m, 2H), 3.31 – 3.22 (m, 7H), 3.15 (t, *J* = 7.2 Hz, 3H), 2.95 – 2.83 (m, 2H), 2.42 – 2.29 (m, 2H), 2.04 – 1.87 (m, 5H). HRMS (ESI) m/z: calcd for C_41_H_43_N_6_O_5_S_2_^+^ [M+H]^+^, 900.3889; found, 900.3894.

**CASPROTAC 2**

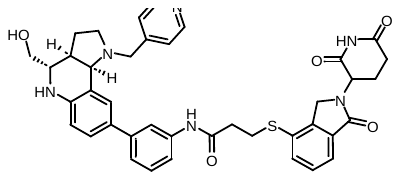

3-((2-(2,6-dioxopiperidin-3-yl)-1-oxoisoindolin-4-yl)thio)-*N*-(3-((3a*R*,4*S*,9b*R*)-4-(hydroxymethyl)-1-(pyridin-4-ylmethyl)-2,3,3a,4,5,9b-hexahydro-1*H*-pyrrolo[3,2-*c*]quinolin-8-yl)phenyl)propanamide (**11b**, 5.9 mg, 31.7% yield, yellow solid). ^1^H NMR (400 MHz, *d_6_-*DMSO) δ 10.97 (s, 1H), 10.05 (s, 1H), 8.70 (d, *J* = 6.0 Hz, 2H), 7.84 (s, 1H), 7.74 – 7.68 (m, 3H), 7.61 – 7.53 (m, 2H), 7.46 – 7.38 (m, 3H), 7.35 – 7.25 (m, 2H), 6.90 (d, *J* = 8.5 Hz, 1H), 5.10 (dd, *J* = 13.3, 5.1 Hz, 1H), 4.82 – 4.75 (m, 1H), 4.67 – 4.53 (m, 1H), 4.35 (d, *J* = 17.7 Hz, 1H), 4.21 (d, *J* = 17.5 Hz, 1H), 3.68 – 3.61 (m, 2H), 3.31 – 3.11 (m, 10H), 2.71 – 2.66 (m, 2H), 2.39 – 2.29 (m, 3H), 2.02 – 1.90 (m, 2H). HRMS (ESI) m/z: calcd for C_40_H_41_N_6_O_5_S_2_^+^ [M+H]^+^, 1034.4468; found, 1034.4461.

**CASPROTAC 3**

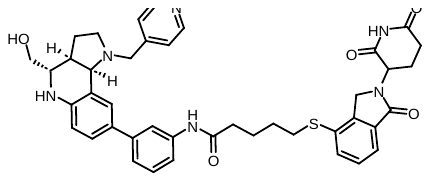

5-((2-(2,6-dioxopiperidin-3-yl)-1-oxoisoindolin-4-yl)thio)-*N*-(3-((3a*R*,4*S*,9b*R*)-4-(hydroxymethyl)-1-(pyridin-4-ylmethyl)-2,3,3a,4,5,9b-hexahydro-1*H*-pyrrolo[3,2-*c*]quinolin-8-yl)phenyl)pentanamide (**11d**, 5.4 mg, 27.9% yield, yellow solid). ^1^H NMR (400 MHz, *d_6_-*DMSO) δ 10.98 (s, 1H), 9.93 (s, 1H), 8.66 (s, 2H), 7.85 (s, 1H), 7.66 – 7.58 (m, 3H), 7.55 (d, *J* = 6.7 Hz, 1H), 7.51 – 7.36 (m, 4H), 7.33 – 7.23 (m, 2H), 6.94 – 6.87 (m, 1H), 5.11 (dd, *J* = 13.2, 5.1 Hz, 1H), 4.82 – 4.50 (m, 2H), 4.35 (d, *J* = 17.5 Hz, 1H), 4.21 (d, *J* = 17.5 Hz, 1H), 3.68 – 3.61 (m, 1H), 3.54 – 3.47 (m, 1H), 3.20 – 3.08 (m, 7H), 2.95 – 2.83 (m, 3H), 2.69 – 2.64 (m, 2H), 2.37 – 2.30 (m, 3H), 2.08 – 1.93 (m, 2H), 1.82 – 1.60 (m, 4H). HRMS (ESI) m/z: calcd for C_42_H_45_N_6_O_5_S^+^ [M+H]^+^, 892.4226; found, 892.4218.

**CASPROTAC 4**

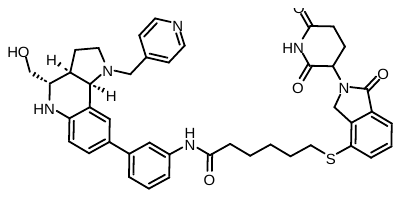

6-((2-(2,6-dioxopiperidin-3-yl)-1-oxoisoindolin-4-yl)thio)-*N*-(3-((3a*R*,4*S*,9b*R*)-4-(hydroxymethyl)-1-(pyridin-4-ylmethyl)-2,3,3a,4,5,9b-hexahydro-1*H*-pyrrolo[3,2-*c*]quinolin-8-yl)phenyl)hexanamide (**11e**, 7.1 mg, 36.0% yield, yellow solid). ^1^H NMR (400 MHz, *d_6_-*DMSO) δ 10.98 (s, 1H), 9.91 (s, 1H), 8.69 (d, *J* = 5.8 Hz, 2H), 7.86 (s, 1H), 7.69 (d, *J* = 5.2 Hz, 2H), 7.62 (d, *J* = 8.5 Hz, 1H), 7.57 – 7.48 (m, 2H), 7.47 – 7.39 (m, 3H), 7.32 – 7.23 (m, 2H), 6.90 (d, *J* = 8.5 Hz, 1H), 5.12 (dd, *J* = 13.3, 5.2 Hz, 1H), 4.83 – 4.58 (m, 2H), 4.34 (d, *J* = 17.4 Hz, 1H), 4.20 (d, *J* = 17.4 Hz, 1H), 3.68 – 3.61 (m, 1H), 3.54 – 3.50 (m, 1H), 3.23 – 3.03 (m, 7H), 2.98 – 2.82 (m, 2H), 2.71 – 2.56 (m, 3H), 2.38 – 2.29 (m, 4H), 2.09 – 1.90 (m, 2H), 1.72 – 1.55 (m, 5H), 1.55 – 1.40 (m, 2H). HRMS (ESI) m/z: calcd for C_43_H_47_N_6_O_5_S^+^ [M+H]^+^, 906.4383; found, 906.4383.

**CASPROTAC 5**

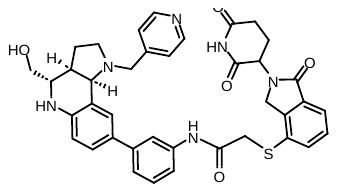

2-((2-(2,6-dioxopiperidin-3-yl)-1-oxoisoindolin-4-yl)thio)-*N*-(3-((3a*R*,4*S*,9b*R*)-4-(hydroxymethyl)-1-(pyridin-4-ylmethyl)-2,3,3a,4,5,9b-hexahydro-1*H*-pyrrolo[3,2-*c*]quinolin-8-yl)phenyl)acetamide (**11a**, 5.2 mg, 28.5% yield, yellow solid). ^1^H NMR (400 MHz, CH_3_OD) δ 8.81 (s, 2H), 8.20 (s, 2H), 7.82 (d, *J* = 7.6 Hz, 2H), 7.69 – 7.60 (m, 1H), 7.55 – 7.40 (m, 3H), 7.36 – 7.28 (m, 2H), 7.24 – 7.14 (m, 1H), 6.89 (d, *J* = 8.4 Hz, 1H), 5.23 – 5.01 (m, 2H), 4.50 (s, 2H), 3.84 (s, 2H), 3.83 – 3.75 (m, 1H), 3.67 – 3.32 (m, 5H), 2.92 – 2.64 (m, 4H), 2.61 – 2.49 (m, 1H), 2.43 – 2.13 (m, 2H), 2.06 – 1.92 (m, 1H). HRMS (ESI) m/z: calcd for C_39_H_39_N_6_O_5_S^+^ [M+H]^+^, 990.4206; found, 990.4206.

**CASPROTAC 6**

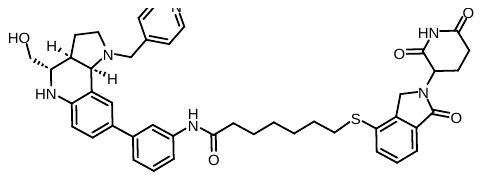

7-((2-(2,6-dioxopiperidin-3-yl)-1-oxoisoindolin-4-yl)thio)-*N*-(3-((3a*R*,4*S*,9b*R*)-4-(hydroxymethyl)-1-(pyridin-4-ylmethyl)-2,3,3a,4,5,9b-hexahydro-1*H*-pyrrolo[3,2-*c*]quinolin-8-yl)phenyl)heptanamide (**11f/GT-03391**, 7.4 mg, 36.8% yield, yellow solid). ^1^H NMR (400 MHz, *d_6_-*DMSO) δ 10.99 (s, 1H), 9.96 (s, 1H), 8.75 (d, *J* = 6.0 Hz, 2H), 7.97 – 7.79 (m, 3H), 7.65 – 7.37 (m, 6H), 7.34 – 7.16 (m, 2H), 6.90 (d, *J* = 8.2 Hz, 1H), 5.13 (dd, *J* = 13.4, 5.1 Hz, 1H), 4.82 (s, 1H), 4.65 (s, 1H), 4.37 – 4.13 (m, 2H), 3.66 (d, *J* = 11.3 Hz, 4H), 3.26 (s, 4H), 3.08 (d, *J* = 7.3 Hz, 2H), 2.88 (s, 1H), 2.59 (d, *J* = 18.5 Hz, 1H), 2.37 – 2.29 (m, 3H), 1.99 (s, 2H), 1.60 (d, *J* = 7.0 Hz, 4H), 1.44 (s, 2H), 1.34 (d, *J* = 7.2 Hz, 2H). ^13^C NMR (126 MHz, *d_6_*-DMSO) δ 172.85, 172.62, 172.45, 170.20, 168.91, 159.79, 157.92, 152.24, 147.62, 140.13, 138.12, 137.73, 137.18, 135.67, 131.94, 130.83, 129.84, 129.11, 128.65, 128.42, 128.28, 127.92, 127.32, 126.29, 123.65, 112.18, 112.11, 110.12, 109.91, 101.20, 100.99, 69.34, 59.17, 56.82, 56.77, 52.67, 50.16, 42.33, 42.12, 38.42, 35.67, 35.33, 34.04, 32.94, 32.61, 29.21, 26.84, 25.92, 25.38, 25.14, 16.16. HRMS (ESI) m/z: calcd for C_44_H_49_N_6_O_5_S^+^ [M+H]^+^, 942.4359; found 942.4367.

**CASPROTAC 7**

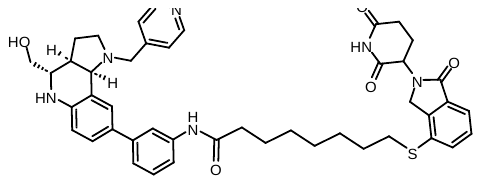

8-((2-(2,6-dioxopiperidin-3-yl)-1-oxoisoindolin-4-yl)thio)-*N*-(3-((3a*R*,4*S*,9b*R*)-4-(hydroxymethyl)-1-(pyridin-4-ylmethyl)-2,3,3a,4,5,9b-hexahydro-1*H*-pyrrolo[3,2-*c*]quinolin-8-yl)phenyl)octanamide (**11g**, 6.8 mg, 33.2% yield, yellow solid). ^1^H NMR (400 MHz, CH_3_OD) δ 8.78 (s, 2H), 8.06 – 7.85 (m, 3H), 7.58 (t, *J* = 7.2 Hz, 2H), 7.53 – 7.42 (m, 3H), 7.38 – 7.21 (m, 3H), 6.89 (d, *J* = 8.8 Hz, 1H), 5.13 (dd, *J* = 13.2, 2.9 Hz, 1H), 5.07 – 4.98 (m, 1H), 4.46 – 4.31 (m, 2H), 3.78 (d, *J* = 8.1 Hz, 1H), 3.62 (dd, *J* = 11.1, 6.3 Hz, 1H), 3.58 – 3.39 (m, 2H), 3.04 (t, *J* = 7.1 Hz, 2H), 2.93 – 2.81 (m, 1H), 2.80 – 2.70 (m, 2H), 2.59 – 2.43 (m, 2H), 2.39 (t, *J* = 7.3 Hz, 2H), 2.27 – 2.09 (m, 2H), 1.73 – 1.61 (m, 4H), 1.51 – 1.37 (m, 6H). HRMS (ESI) m/z: calcd for C_45_H_51_N_6_O_5_S^+^ [M+H]^+^, 956.4515; found, 956.4524.

**CASPROTAC 8**

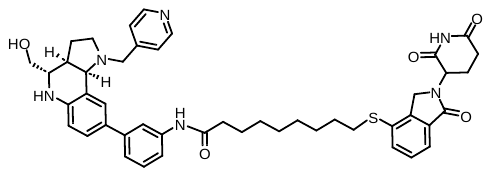

9-((2-(2,6-dioxopiperidin-3-yl)-1-oxoisoindolin-4-yl)thio)-*N*-(3-((3a*R*,4*S*,9b*R*)-4-(hydroxymethyl)-1-(pyridin-4-ylmethyl)-2,3,3a,4,5,9b-hexahydro-1*H*-pyrrolo[3,2-*c*]quinolin-8-yl)phenyl)nonanamide (**11h**, 7.9 mg, 37.9% yield, yellow solid). ^1^H NMR (400 MHz, CH_3_OD) δ 8.75 (s, 2H), 8.02 – 7.84 (m, 3H), 7.60 (d, *J* = 7.6 Hz, 2H), 7.54 – 7.43 (m, 3H), 7.38 – 7.21 (m, 3H), 6.88 (d, *J* = 8.3 Hz, 1H), 5.14 (dd, *J* = 13.4, 5.0 Hz, 1H), 5.04 – 4.96 (m, 1H), 4.47 – 4.32 (m, 2H), 3.83 – 3.73 (m, 1H), 3.65 – 3.57 (m, 1H), 3.53 – 3.36 (m, 2H), 3.02 (t, *J* = 7.0 Hz, 2H), 2.94 – 2.82 (m, 1H), 2.81 – 2.68 (m, 2H), 2.58 – 2.46 (m, 2H), 2.38 (t, *J* = 7.3 Hz, 2H), 2.25 – 2.11 (m, 2H), 1.77 – 1.59 (m, 4H), 1.50 – 1.33 (m, 8H). HRMS (ESI) m/z: calcd for C_46_H_53_N_6_O_5_S^+^ [M+H]^+^, 970.4672; found, 970.4668.

**CASPROTAC 9**

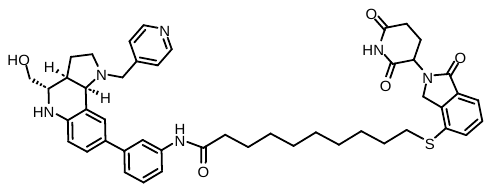

10-((2-(2,6-dioxopiperidin-3-yl)-1-oxoisoindolin-4-yl)thio)-*N*-(3-((3a*R*,4*S*,9b*R*)-4-(hydroxymethyl)-1-(pyridin-4-ylmethyl)-2,3,3a,4,5,9b-hexahydro-1*H*-pyrrolo[3,2-*c*]quinolin-8-yl)phenyl)decanamide (**11i**, 8.3 mg, 39.2% yield, yellow solid). ^1^H NMR (400 MHz, *d_6_-*DMSO) δ NMR (400 MHz, MeOD) δ 8.78 – 8.63 (m, 2H), 8.04 – 7.83 (m, 3H), 7.64 – 7.43 (m, 5H), 7.35 – 7.20 (m, 3H), 6.88 (d, *J* = 8.6 Hz, 1H), 5.18 – 5.09 (m, 1H), 5.05 – 4.97 (m, 1H), 4.46 – 4.33 (m, 2H), 3.82 – 3.76 (m, 1H), 3.62 (dd, *J* = 11.2, 6.4 Hz, 1H), 3.54 – 3.38 (m, 2H), 3.01 (t, *J* = 7.0 Hz, 2H), 2.91 – 2.81 (m, 1H), 2.80 – 2.68 (m, 2H), 2.59 – 2.47 (m, 2H), 2.39 (t, *J* = 7.4 Hz, 2H), 2.26 – 2.11 (m, 2H), 1.76 – 1.57 (m, 4H), 1.37 – 1.27 (m, 10H). HRMS (ESI) m/z: calcd for C_47_H_55_N_6_O_5_S^+^ [M+H]^+^, 984.4828; found, 984.4834.

**CASPROTAC 10**

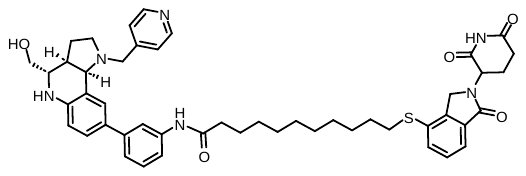

11-((2-(2,6-dioxopiperidin-3-yl)-1-oxoisoindolin-4-yl)thio)-*N*-(3-((3a*R*,4*S*,9b*R*)-4-(hydroxymethyl)-1-(pyridin-4-ylmethyl)-2,3,3a,4,5,9b-hexahydro-1*H*-pyrrolo[3,2-*c*]quinolin-8-yl)phenyl)undecanamide (**11j**, 8.6 mg, 39.9% yield, yellow solid). ^1^H NMR (400 MHz, CH_3_OD) δ 8.69 (s, 2H), 7.96 (s, 1H), 7.86 – 7.73 (m, 2H), 7.66 – 7.57 (m, 2H), 7.52 – 7.44 (m, 3H), 7.37 – 7.21 (m, 3H), 6.88 (d, *J* = 7.6 Hz, 1H), 5.19 – 5.09 (m, 1H), 5.01 – 4.96 (m, 1H), 4.46 – 4.37 (m, 2H), 3.81 – 3.73 (m, 1H), 3.65 – 3.59 (m, 1H), 3.50 – 3.44 (m, 2H), 3.02 (t, *J* = 7.4 Hz, 2H), 2.90 – 2.82 (m, 1H), 2.79 – 2.68 (m, 2H), 2.55 – 2.44 (m, 2H), 2.39 (t, *J* = 7.4 Hz, 2H), 2.22 – 2.13 (m, 2H), 1.79 – 1.55 (m, 6H), 1.34 – 1.23 (m, 10H). HRMS (ESI) m/z: calcd for C_48_H_57_N_6_O_5_S^+^ [M+H]^+^, 984.4828; found, 984.4834.
