## Supplementary figures for "PROTAC molecule-mediated SpCas9 protein degradation for precise genome editing"

### Slide 1
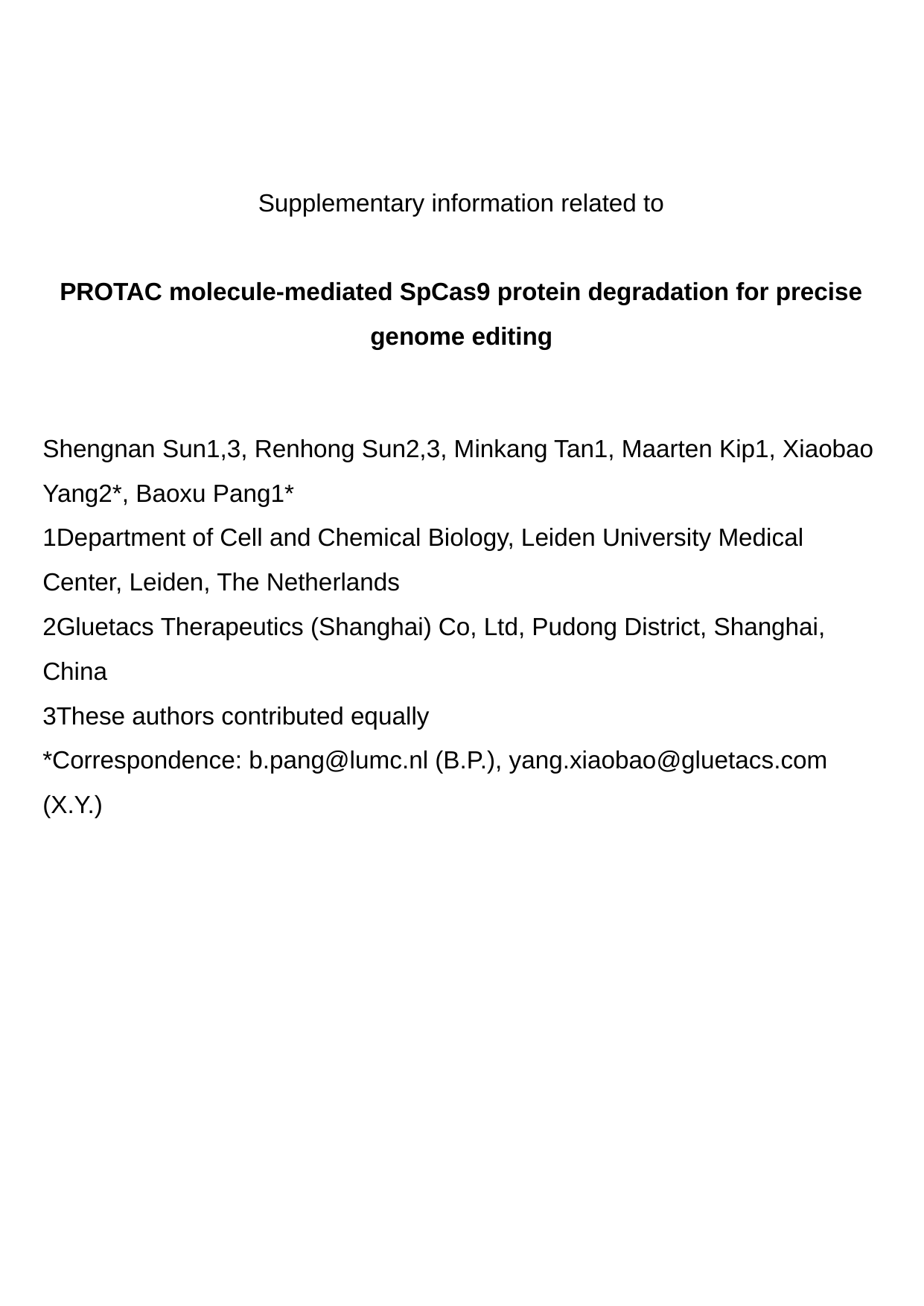

Supplementary information related to
PROTAC molecule-mediated SpCas9 protein degradation for precise genome editing
#### Shengnan Sun1,3, Renhong Sun2,3, Minkang Tan1, Maarten Kip1, Xiaobao Yang2*, Baoxu Pang1* 1Department of Cell and Chemical Biology, Leiden University Medical Center, Leiden, The Netherlands 2Gluetacs Therapeutics (Shanghai) Co, Ltd, Pudong District, Shanghai, China 3These authors contributed equally *Correspondence: (B.P.), (X.Y.)

### Slide 2
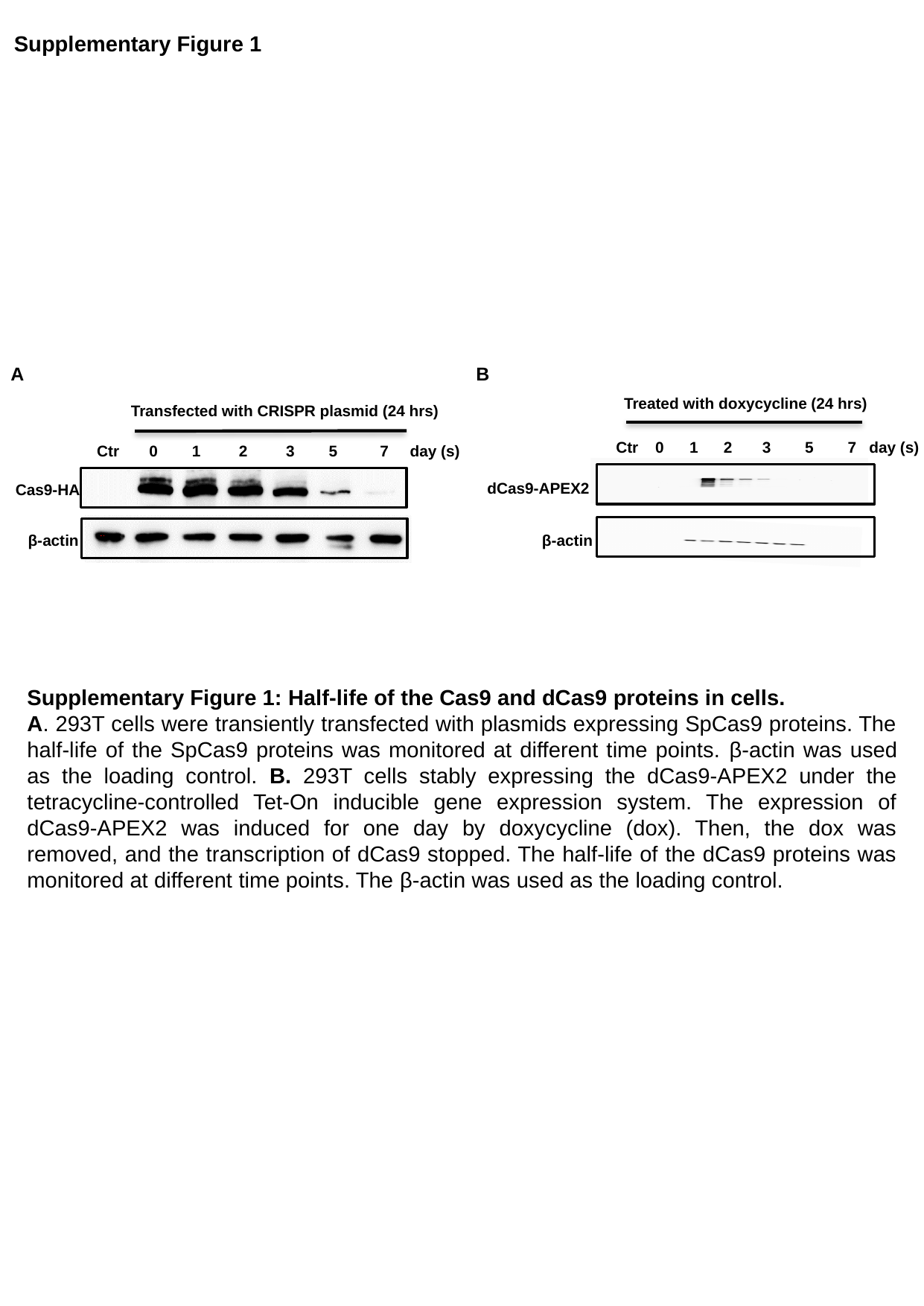

Supplementary Figure 1
A
B
Treated with doxycycline (24 hrs)
Transfected with CRISPR plasmid (24 hrs)
 Ctr 0 1 2 3 5 7 day (s)
 Ctr 0 1 2 3 5 7 day (s)
dCas9-APEX2
Cas9-HA
β-actin
β-actin
Supplementary Figure 1: Half-life of the Cas9 and dCas9 proteins in cells.
A. 293T cells were transiently transfected with plasmids expressing SpCas9 proteins. The half-life of the SpCas9 proteins was monitored at different time points. β-actin was used as the loading control. B. 293T cells stably expressing the dCas9-APEX2 under the tetracycline-controlled Tet-On inducible gene expression system. The expression of dCas9-APEX2 was induced for one day by doxycycline (dox). Then, the dox was removed, and the transcription of dCas9 stopped. The half-life of the dCas9 proteins was monitored at different time points. The β-actin was used as the loading control.

### Slide 3
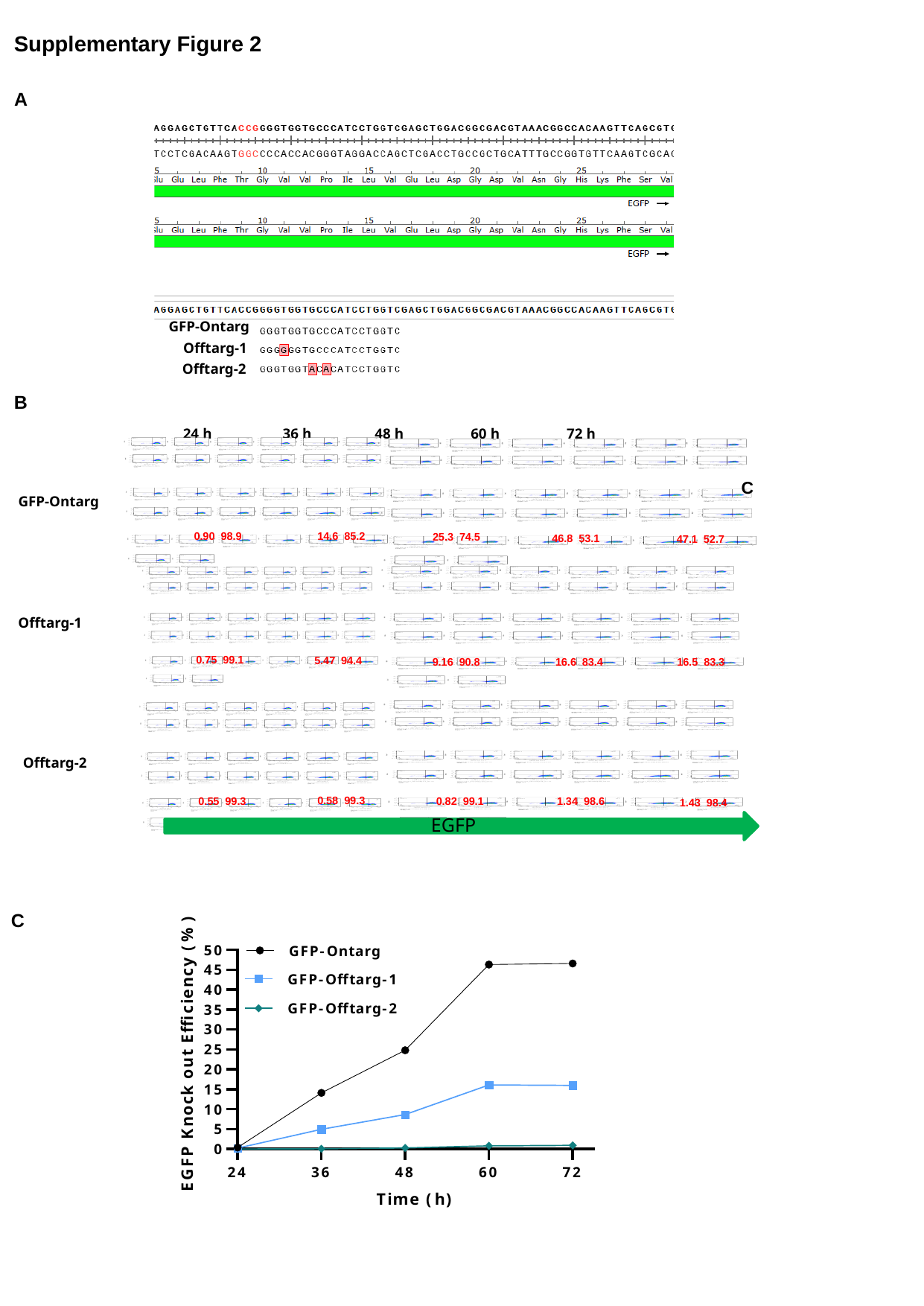

Supplementary Figure 2
A
GFP-Ontarg
 Offtarg-1
 Offtarg-2
B
 24 h 36 h 48 h 60 h 72 h
 0.90 98.9
 14.6 85.2
 25.3 74.5
 46.8 53.1
 47.1 52.7
 0.75 99.1
 5.47 94.4
 16.5 83.3
 9.16 90.8
 16.6 83.4
 0.58 99.3
 1.34 98.6
 0.82 99.1
 0.55 99.3
 1.43 98.4
GFP-Ontarg
 Offtarg-1
 Offtarg-2
EGFP
C
C

### Slide 4
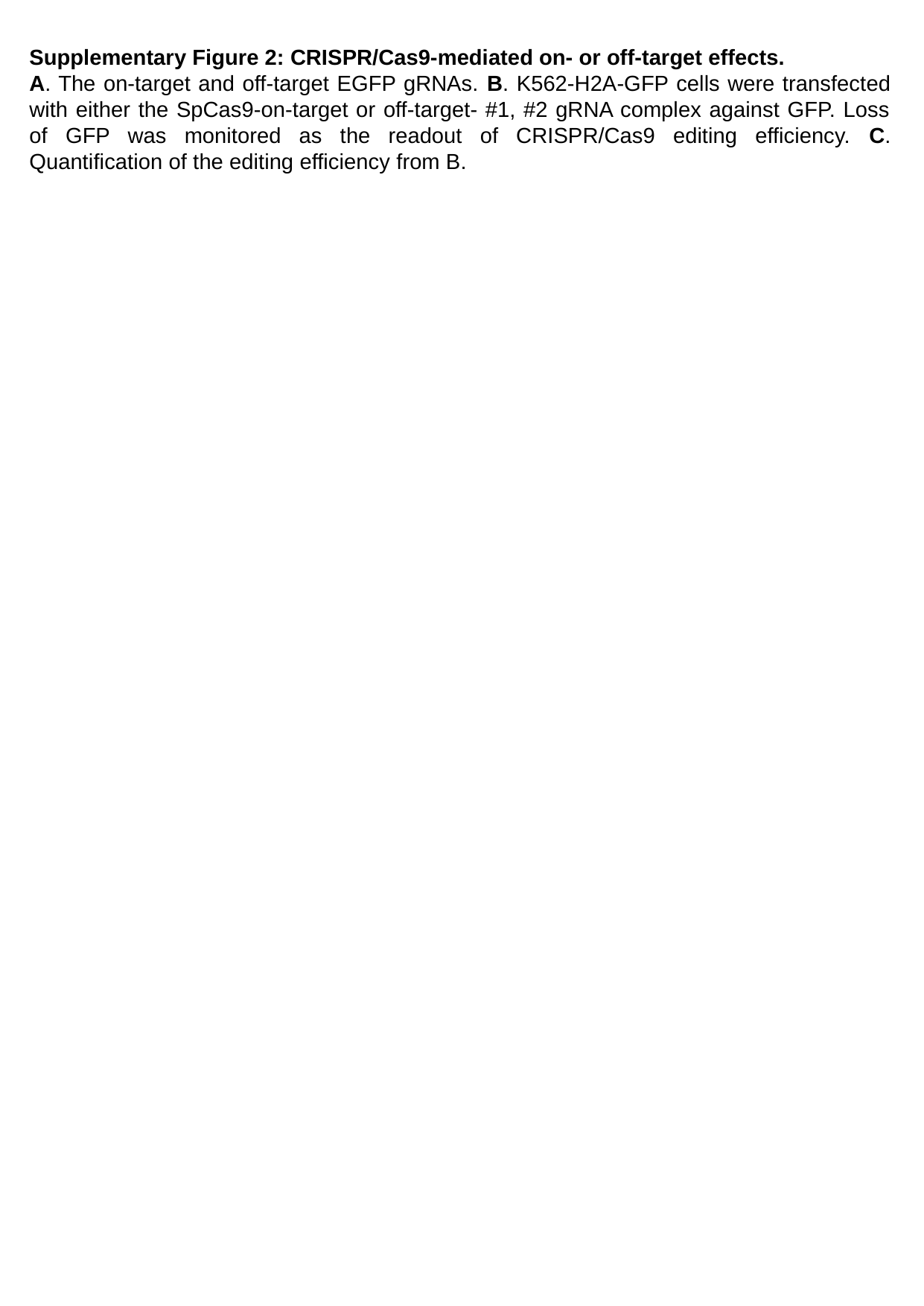

Supplementary Figure 2: CRISPR/Cas9-mediated on- or off-target effects.
A. The on-target and off-target EGFP gRNAs. B. K562-H2A-GFP cells were transfected with either the SpCas9-on-target or off-target- #1, #2 gRNA complex against GFP. Loss of GFP was monitored as the readout of CRISPR/Cas9 editing efficiency. C. Quantification of the editing efficiency from B.
